# Psychedelics Restructure the Global Workspace of Consciousness

**DOI:** 10.64898/2026.09.28.755172

**Authors:** Parsa Yousefi, Kenneth Shinozuka, Patrick Vizeli, Friederike Holze, Anna M. Becker, Felix Müller, Matthias E. Liechti, Gregor Hasler, Carolyn I. Rodríguez, Andrea Luppi, Pedro A.M. Mediano

## Abstract

Psychoactive drugs profoundly alter consciousness, but the neural mechanisms by which they do so remain poorly understood. Global workspace theory may provide a framework for investigating these mechanisms. The global workspace, a candidate neural substrate of consciousness, is a set of brain regions that integrate and disseminate information to the rest of the brain. According to a recent model, the integrators, or “gateways,” are characterized by synergistic interactions with other brain regions, while the disseminators, or “broadcasters,” exhibit redundant interactions. Here, we measured the effects of various psychoactive drugs, including “classic” psychedelics like LSD, mescaline, and psilocybin, “atypical” psychedelics (entactogens) like MDMA, and the stimulant D-amphetamine, on the global workspace. In a secondary analysis of resting-state functional magnetic resonance imaging (fMRI) data on all these drugs, we found that classic psychedelics are associated with significant increases in the number of gateways in the frontoparietal control and dorsal attention networks. Meanwhile, MDMA and amphetamine reduced the number of gateways in the visual and somatomotor networks. The regional topographies of both synergy and redundancy distinguished classic psychedelics from other drugs. Finally, drugs producing similar changes in synergy and redundancy also produced similar profiles of subjective effects. Overall, our findings provide the first direct evidence that psychedelics may alter the synergistic and redundant interactions that characterize the global workspace of consciousness.

## INTRODUCTION

Psychedelics are potentially effective treatments for a wide range of psychiatric and neurological conditions, including depression, anxiety, post-traumatic stress disorder, substance use disorders, and possibly brain injury.^1–10^ They are typically separated into two classes: “classic” psychedelics like lysergic acid diethylamide (LSD), psilocybin, and mescaline, which are agonists at the 5-HT_2A_ (serotonin-2A) receptor, and “atypical” psychedelics like 3,4-methylenedioxymethamphetamine (MDMA), an entactogen/empathogen that acts as monoamine and oxytocin releaser.^11^

Classic psychedelics induce strongly altered states of consciousness, whose “phenomenology,” or subjective experience, includes ego dissolution (loss of sense of self), euphoria, and a sense of unity with the environment.^12,13^ MDMA, by contrast, produces a milder alteration of consciousness, mainly leading to euphoria, emotional openness, and feelings of closeness to others, with little ego dissolution and fewer perceptual hallucinations.^14^ Several neuroimaging studies have measured correlations between these experiences and brain activity, for instance finding that ego dissolution is associated with global functional connectivity (FC) and decreases in the activity and FC of specific networks like the default mode network (DMN).^15–29^ Meanwhile, acute MDMA produces distributed changes in fMRI measures of brain function, including reduced activity or perfusion in medial temporal regions and altered functional connectivity involving the amygdala, hippocampus, medial prefrontal cortex, and insula, while its effects on large-scale between-network connectivity appear relatively modest.^30–32^ Computational modeling has extended this work by showing how psychedelics reshape whole-brain dynamics, including changes in dynamical flexibility, metastability, and the brain’s repertoire of accessible states.^33–35^ Yet these models have generally focused on explaining the neural effects of psychedelics themselves, rather than how those neural changes produce particular alterations in conscious experience. Thus, despite important advances in modeling psychedelic brain states, there remains limited model-based evidence directly linking psychedelic-induced changes in brain dynamics to changes in consciousness.

One framework that may help bridge this gap is Global Workspace Theory. This theory proposes that the brain contains many specialized systems that process specific kinds of information largely in parallel (e.g., the visual cortex processes visual information), but that information becomes consciously accessible only when it enters a shared “workspace” and is broadcast across the brain.^36,37^ The neuronal formulation of this theory, Global Neuronal Workspace Theory, proposes that this transition occurs through nonlinear “ignition,” whereby recurrent interactions amplify and sustain an initially local neural representation across distributed cortical regions, making that information globally available to the rest of the brain.^38–41^ Once information enters this workspace, it becomes available to otherwise specialized systems involved in processes such as memory, evaluation, decision-making, and action. This framework therefore links consciousness to specific properties of large-scale information flow rather than to activity within any single brain region or network.

More recent work has begun to formalize the global workspace using information theory. In particular, Luppi and colleagues proposed a “*synergistic* global workspace,” whereby workspace regions are identified as those with the highest capacity to integrate information across distant connections.^42^ This capacity is diminished during anesthesia and disorders of consciousness and restored during recovery. Based on integrated information decomposition (ΦID), “gateways” are defined as regions in the global workspace that gather and integrate information from specialized brain modules, while “broadcasters” are regions that redistribute this information throughout the brain.^42–44^ These roles are characterized by high synergistic and redundant participation coefficients (PCs), respectively: synergy reflects information that becomes available only when the activity of multiple regions is considered jointly, whereas redundancy reflects information that is shared across regions and can therefore be obtained from either region alone. Although gateways are a recently developed construct whose test– retest reliability has not yet been systematically established, their anatomical distribution is robust to alternative parcellations and network definitions, and gateway-mediated information integration is selectively disrupted during both pharmacological and pathological loss of consciousness and restored following recovery, providing initial evidence for their biological and clinical relevance.^42^

Whereas the conventional ignition framework identifies the neural event that causes information to become globally available, the ΦID model characterizes the complementary operations through which information is integrated into and distributed across the workspace. It therefore provides a suitable framework for testing how psychedelics reorganize these operations.

Here, we measured the synergistic global workspace in two previously acquired resting-state fMRI datasets, both collected on the same scanner at the University of Basel. In the first, LSD, amphetamine, and MDMA were administered to a group of healthy volunteers in a within-subjects, crossover design (LAM study).^14^ In the second, LSD, psilocybin, and mescaline were administered to a distinct group of healthy volunteers, also in a within-subjects, crossover design (LPM study).^45^ We hypothesized that the classic psychedelics would increase the number of gateways in the global workspace, based on prior evidence that these drugs increase the diversity of brain dynamics and weaken the segregation between normally distinct functional networks.^11,16,21,26,27,29,32,46–48^ Because gateways and broadcasters are primarily located in the DMN and FPN, respectively,^42^ our primary outcome measures were the changes in the roles of these two networks. We also hypothesized that the drugs could be separated into their respective pharmacological classes based on their synergy and redundancy PCs.

## RESULTS

### Classic psychedelics increase the number of gateways in the frontoparietal control network and dorsal attention network

An overview of the study design is shown in **Figure 1**. Using data from the Human Connectome Project (HCP),^49^ we applied the global-workspace definition described by Luppi et al. (2024).^42^ The majority of the regions in the subcortex, frontoparietal network (FPN), default mode network (DMN), and limbic network (LIM) were included in the 235-region global workspace (**Supplementary Figure 1**; note the entire brain parcellation contains 454 regions).

**Figure 1.**
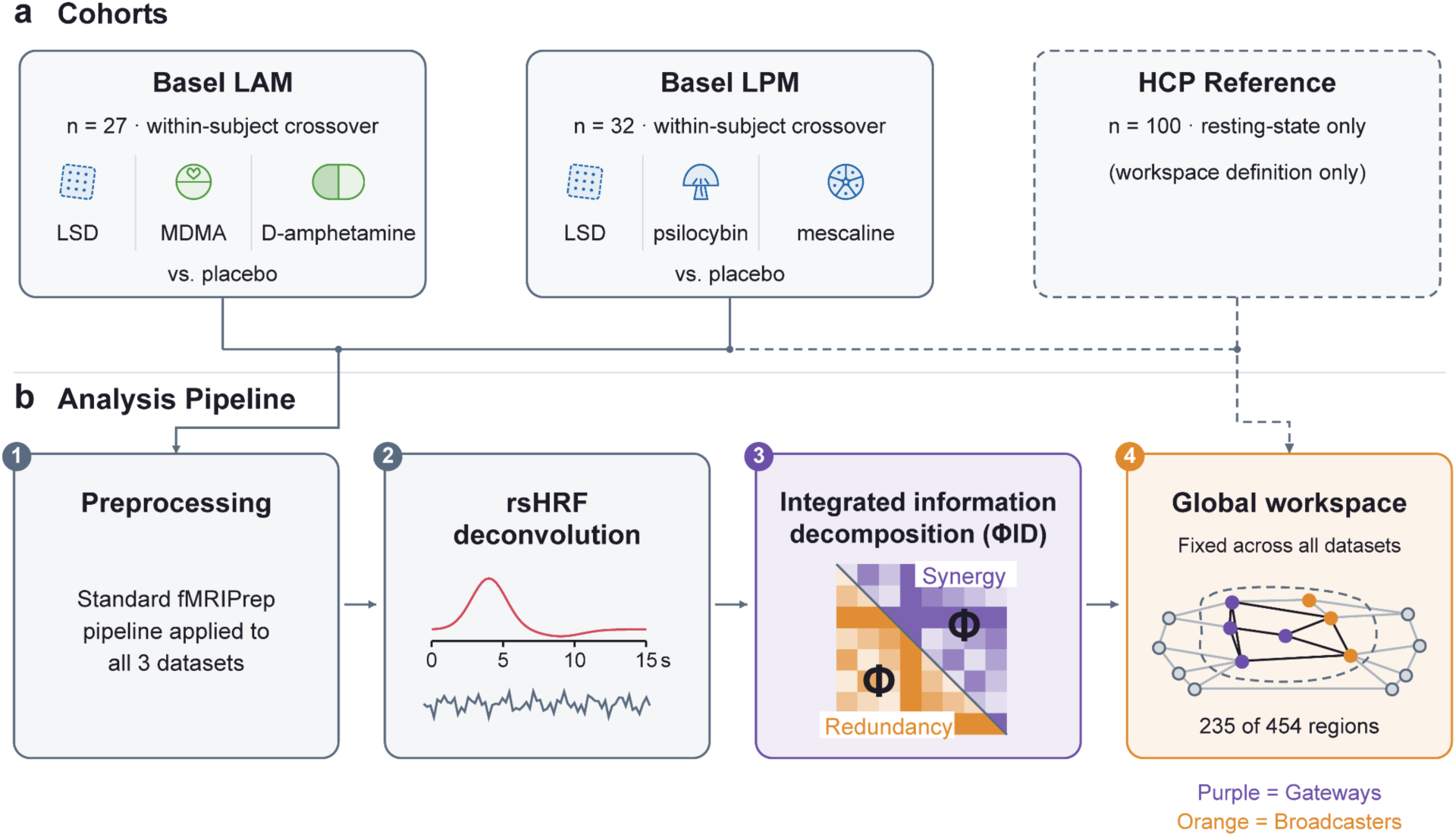
Overview. Using integrated information decomposition, the global workspace of consciousness was measured in six active drug conditions. (a) fMRI data were analyzed from three cohorts: the HCP reference sample (n = 100), used only to define the global workspace; the LAM study (*n* = 27), in which LSD, MDMA, D-amphetamine, and placebo were administered to healthy participants in a within-subjects crossover;^14^ and the Basel LPM study (*n* = 32), in which LSD, psilocybin, mescaline, and placebo were administered to healthy participants in a within-subjects crossover.^45^ (b) Raw fMRI data were obtained from all three studies. An identical preprocessing pipeline, consisting of fMRIPrep, CONN denoising, and parcellation into 454 regions (Schaefer-400 cortical + Tian-54 subcortical), and rsHRF deconvolution, was applied to all three datasets. Integrated information decomposition was used to determine synergistic and redundant interactions between pairs of regions. The global workspace comprises 235 regions whose synergy-strength rank exceeds their redundancy-strength rank in the HCP reference data. Within this workspace, gateways have a higher rank in synergy participation coefficient than in redundancy participation coefficient; broadcasters show the reverse rank relationship. Equal ranks are assigned to neither role.

Psychoactive drugs differentially altered the number of gateways in specific networks (**Figure 2**; **Supplementary Table 2**; the complete distribution across networks and drug conditions is shown in **Supplementary Figure 2**). Note that an increase in gateways in a given network generally implies an equivalent decrease in broadcasters. Across the four classic-psychedelic conditions, the number of FPN gateways increased by an average of 4.37 regions. This increase was individually significant under LSD (LAM; +4.35 regions, *q* < .001), LSD (LPM; +4.60 regions, *q* < .001), psilocybin (+4.52 regions, *q* < .001), and mescaline (+4.03 regions, *q* = .002). DMN gateway counts increased by an average of 1.32 regions across the four conditions, although the increase was individually significant only under LSD (LPM; +1.47 regions, *q* = .032) and mescaline (+1.81 regions, *q* = .035). Among the other networks, the number of gateways in the dorsal attention network (DAN) increased significantly under all four classic-psychedelic conditions (*q* = .001–.032). The number of gateways decreased in the visual network (VIS) under both LSD conditions, in the somatomotor network (SMN) under LSD (LAM) and psilocybin, in the LIM under LSD (LPM) and mescaline, and in the ventral attention network (VAN) under psilocybin. MDMA and amphetamine both reduced VIS and SMN gateway counts, while MDMA additionally increased DAN gateway counts. In general, both classic psychedelics and stimulants/entactogens increased gateways in higher-order regions along the cortical hierarchy (**Supplementary Figure 4**).

**Figure 2.**
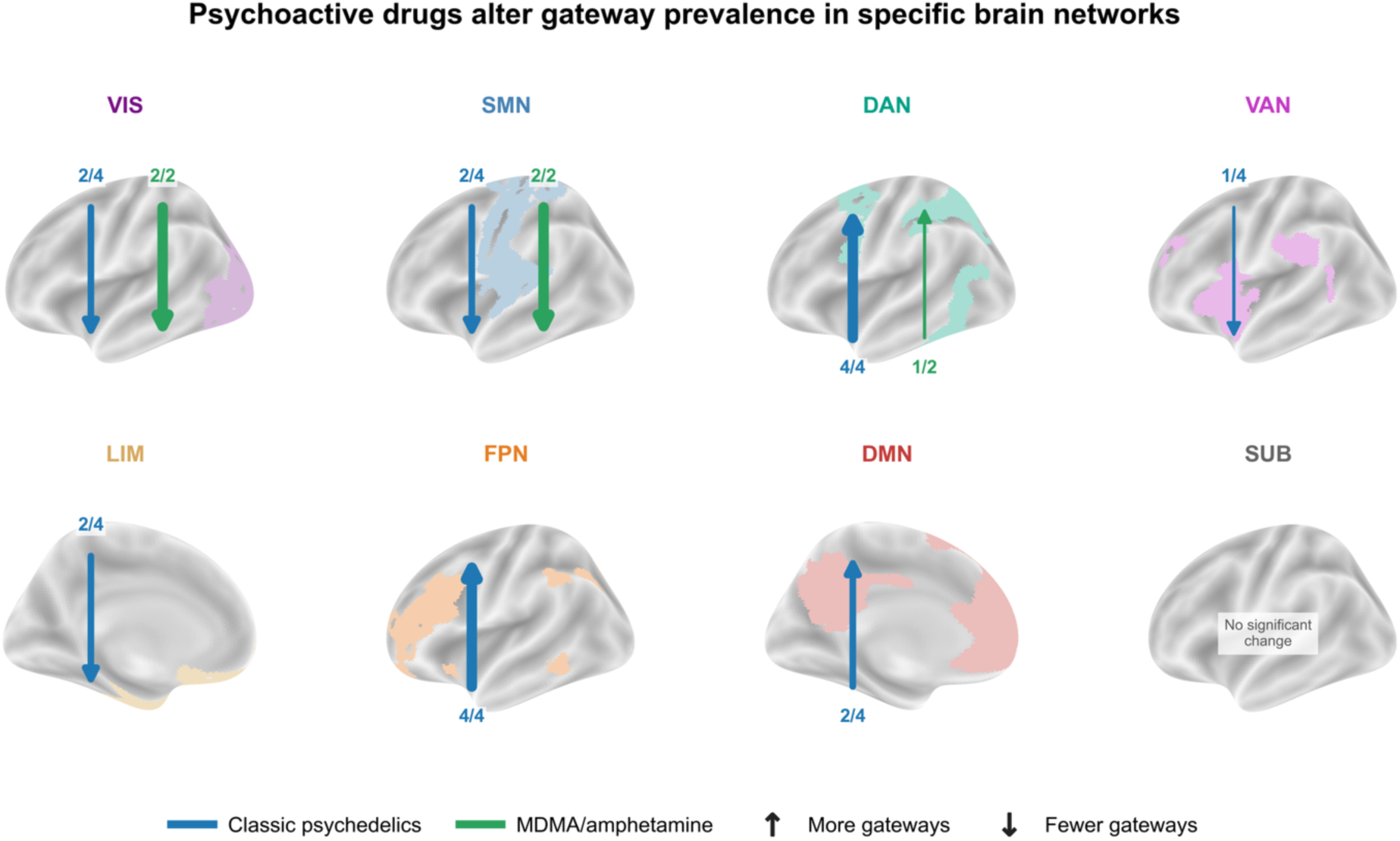
Classic psychedelics increase the number of gateways in certain higher-order networks, while MDMA and amphetamine both decrease the number of gateways in lower-order networks. Brain maps highlight the seven cortical networks of the Schaefer-400 atlas; the Tian-54 subcortical regions were treated as an eighth network. Arrows summarize significant drug-versus-control changes in gateway counts within each network. Blue arrows represent classic psychedelics and green arrows represent MDMA or amphetamine; upward arrows indicate more gateways and downward arrows indicate fewer gateways. Arrow thickness reflects the number of conditions within each class showing the effect, and the accompanying fraction gives the exact number; for example, 4/4 indicates significance in all four classic-psychedelic conditions, yielding a maximally thick arrow. FDR correction was applied separately within each condition across the two primary networks, DMN and FPN, and across the remaining six exploratory networks; correction was not performed across drug conditions. All classic psychedelics increased gateways in the FPN and DAN, while both MDMA and amphetamine reduced gateways in the VIS and SMN. Individual effect sizes and significance levels are shown in Supplementary Figure 3 and Supplementary Table 2.

In addition to changing the number of gateways in specific networks, classic psychedelics also altered the distribution of synergistic and redundant interactions between the global workspace and the rest of the brain. Gateways are regions with relatively high synergy PC, reflecting synergistic interactions distributed broadly across brain networks. Meanwhile, broadcasters are regions with relatively high redundancy PC, indicating redundant interactions with a diverse set of networks. Mean redundancy PC increased in all four classic-psychedelic conditions (Hedges’ *g*_z_ = 1.28-2.05, *q <* .001), as well as under MDMA (*g*_z_ = 0.66, *q* = .002) and amphetamine (*g*_z_ = 1.13, *q* < .001) (**Supplementary Figure 5**). Mean synergy PC increased under LSD in LPM (*g*_z_ = 0.70, *q* = .001), psilocybin (*g*_z_ = 1.05, *q* < .001), and mescaline (*g*_z_ = 0.58, *q* = .005), but not under LSD in LAM or any non-classic psychedelic condition. Thus, broader distribution of redundancy across networks was common to several drugs, whereas increased synergy PC occurred in most but not all classic psychedelics.

None of the correlations between drug-related changes in framewise displacement (FD) and changes in gateway counts survived false discovery rate (FDR) correction across the 48 drug condition-by-network tests. Re-estimating the gateway-count effects while adjusting for FD preserved all significant FPN gateway increases. The DMN gateway increase under LSD (LPM) no longer survived (motion-adjusted *q* = .074), whereas a previously non-significant decrease in LIM gateways under psilocybin became significant (motion-adjusted *q* = .038). All other changes in gateway counts were unchanged after adjusting for head motion (**Supplementary Table 3**). Similarly, none of the correlations between FD and mean synergy-or redundancy-PC changes survived correction across the 12 condition-by-measure tests. Adjusting for FD did not change the statistical significance of any change in mean synergy or redundancy PC effect (**Supplementary Table 4**).

### Synergy and redundancy separate drug classes

We next determined whether the synergy and redundancy PC would be effective at separating the drug classes. Hierarchical clustering provided an initial visualization of the relationships among the six drug conditions (**Supplementary Figure 6**). Only redundancy PC correctly grouped together all four classic-psychedelic conditions into a separate cluster than MDMA and amphetamine, whereas synergy PC clustered LSD (LAM) together with MDMA and amphetamine. The corresponding matrices of between-drug Pearson correlations in synergy and redundancy PC are presented in **Supplementary Figure 7**.

However, this clustering was treated as descriptive because some clusters could reflect study design rather than genuine similarities in global workspace activity. Drug conditions within LAM or LPM shared participants and placebo scans, whereas drug conditions from different studies did not. Consequently, an apparent cluster could arise partly from study-specific acquisition or shared-control effects.

We therefore tested three complementary aspects of drug-class separation using participant-level randomization tests that preserved the within-subject, crossover structure of each study (**Figure 3**). Within LAM, synergy PC and redundancy PC both separated LSD from MDMA and amphetamine: the separation statistic was 0.127 for synergy PC (*q* = .023) and 0.271 for redundancy PC (*q* < .001; **Figure 3a**). LSD from LAM was more similar to the three LPM classic psychedelics than expected under condition-label randomization. Its mean similarity with the LPM psychedelics exceeded the null expectation by 0.179 for synergy PC (*q* = .011) and 0.217 for redundancy PC (*q* < .001; **Figure 3b**). Conversely, the LPM psychedelics were less similar to MDMA and amphetamine than expected, with differences from the null of 0.106 for synergy PC (*q* = .008) and 0.200 for redundancy PC (*q* < .001; **Figure 3c**). Thus, both the synergistic and redundant topographies distinguished classic psychedelics from stimulants/entactogens within LAM and generalized across the independent LAM and LPM cohorts.

**Figure 3.**
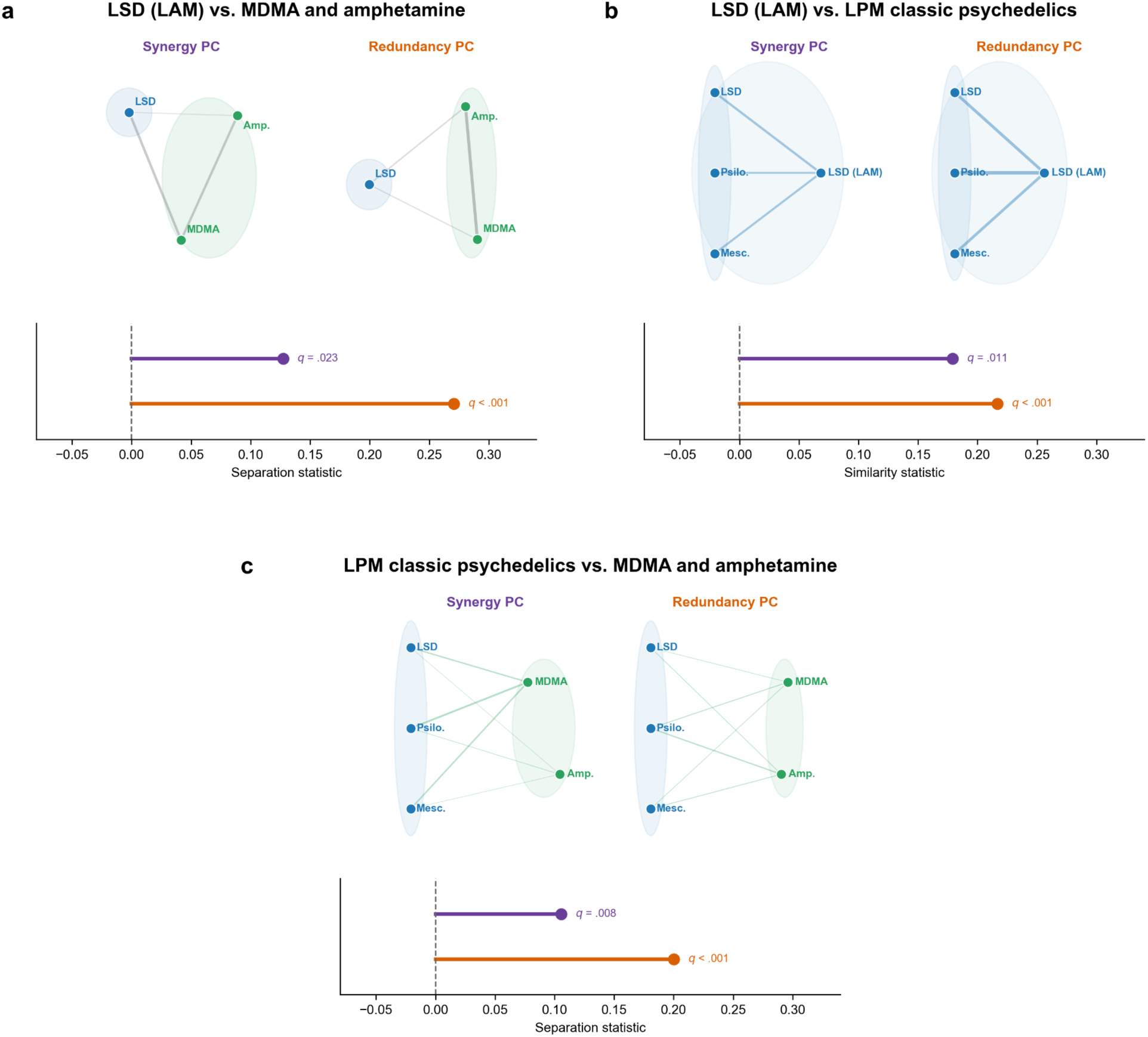
Synergy and redundancy PC separate the drug classes. Differences in synergy and redundancy PC between each drug condition and placebo were represented as Hedges’ *g*_av_ values. Lines represent distances between pairs of drug conditions in terms of synergy or redundancy PC. Thicker lines indicate more similar synergy or redundancy PCs between drug conditions. Blue shading denotes classic psychedelics, and green shading denotes stimulants/entactogens. Purple and orange distinguish analyses based on synergy PC and redundancy PC, respectively. The plots below each graphic show the corresponding separation or similarity statistic relative to the null, represented by the vertical dashed line at zero. (a) Within LAM, both synergy and redundancy PC significantly separated LSD from MDMA and amphetamine. (b) LSD from LAM was significantly similar to the three LPM psychedelics in both synergy and redundancy PC. (c) Both synergy and redundancy PC significantly separated the three LPM psychedelics from MDMA and amphetamine. Drug condition labels were randomized within participants while placebo was held fixed, and the regional effect profiles for synergy and redundancy PC and the corresponding statistics were recomputed across 100,000 randomizations. FDR correction was applied across the two measures in panel a) and separately across the four cross-study tests in panels b) and c). P-values are one-sided in the hypothesized direction: greater separation than the null in (a) and (c), and greater similarity than the null in (b).

Which brain networks contribute to differences in synergy and redundancy PC between drug classes? To address this question, we repeated each classification analysis after removing the regions belonging to one network at a time (**Figure 4**). No single network accounted for the overall separation between classic psychedelics and stimulants/entactogens. Instead, different networks made distinct contributions to the between-drug comparisons. For redundancy PC, removing VIS increased within-LAM class separation and reduced the similarity of LSD (LAM) to the LPM psychedelics; removing SMN reduced within-LAM class separation, and removing DMN increased cross-study psychedelic similarity. Removing VIS also increased the separation of the LPM psychedelics from MDMA and amphetamine for both synergy and redundancy PC. These effects survived the corresponding FDR corrections.

**Figure 4.**
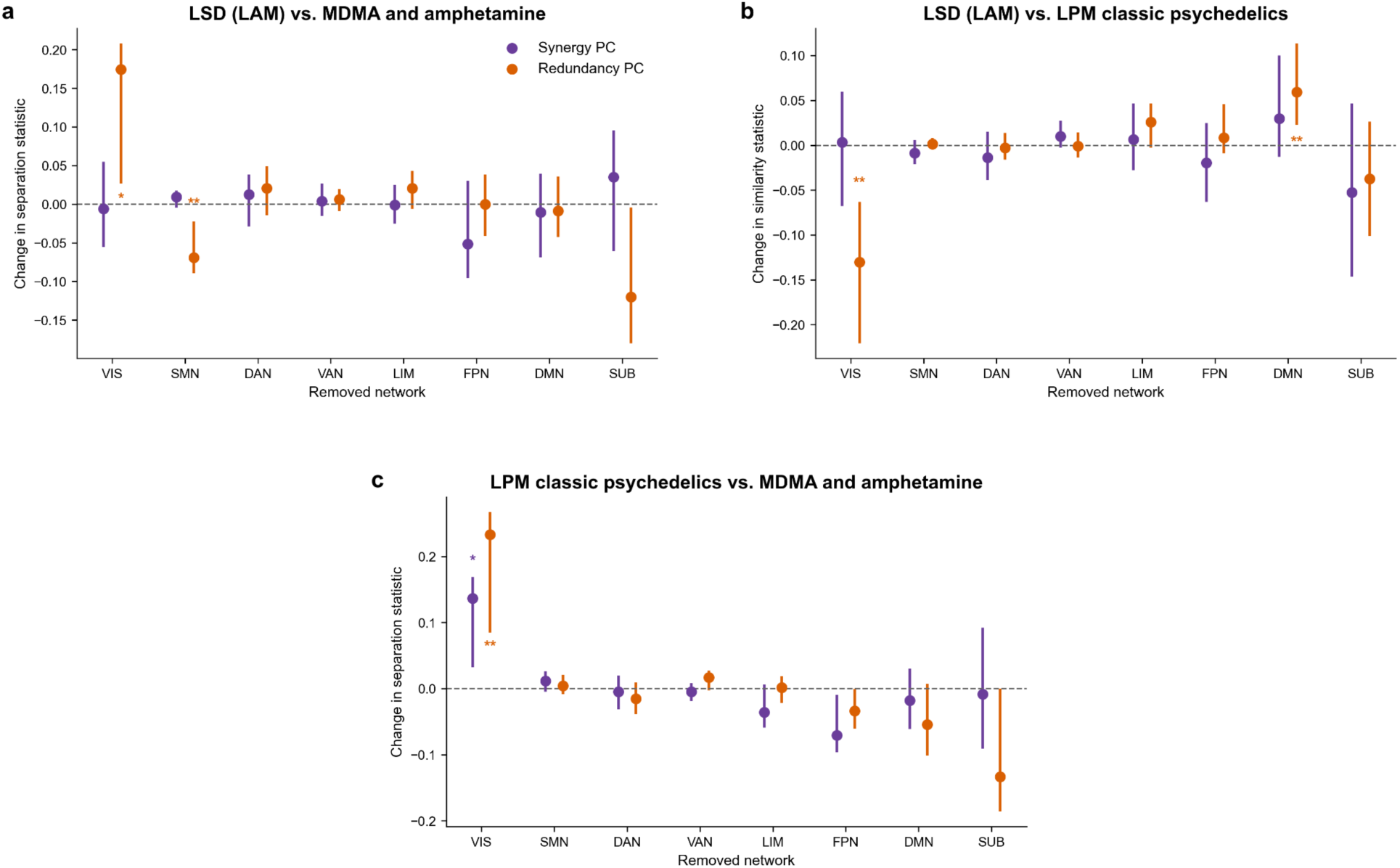
Differences between drugs in synergy and redundancy PC depend on specific networks. The contribution of each network was evaluated based on a leave-one-network-out analysis: the network’s regions were removed from the synergy and redundancy PCs and the relevant classification statistic was recalculated. Positive values indicate that removing the network increased separation or similarity, whereas negative values indicate that its removal reduced the corresponding statistic. (a) Within the LAM study, removal of VIS significantly increased the separation of LSD from MDMA and amphetamine based on redundancy PC, whereas removal of SMN significantly reduced it. (b) Based on redundancy PC, removal of VIS reduced the similarity of LSD (LAM) to the three LPM classic psychedelics, while removal of DMN increased their similarity. (c) Removal of VIS increased the separation of the LPM classic psychedelics from MDMA and amphetamine based on both synergy and redundancy PC. Error bars show participant-bootstrap 95% confidence intervals from 10,000 resamples. Asterisks indicate significant changes after FDR correction across the eight removed networks, performed separately for each PC and classification statistic. * *q* < .05, ** *q* < .01. Bootstrap p-values are percentile-based.

### Synergy and redundancy track phenomenology

Finally, we asked whether the differences in synergy and redundancy between drug classes corresponded to the differences in phenomenology between drug classes. First, the effect sizes of the synergy PC and redundancy PC (the neural “profiles”) were each correlated between drug conditions. Then, the phenomenology “profiles” (ratings of each drug on the 11 lower-order subscales of the 5-Dimensional Altered States of Consciousness [5D-ASC] questionnaire,^50^ which were acquired in the LAM and LPM studies; see **Supplementary Table 6**) were each correlated between drug conditions. We measured the similarities of the between-drug correlations in the neural profiles with the between-drug correlations in the phenomenology profiles (**Figure 5**). The synergy PC and phenomenology profile had very similar between-drug correlations (ρ = .92, *q* = .003). The same was true of the redundancy PC (ρ = .76, *q* = .029). Furthermore, between-drug differences in synergy PC were significantly associated with between-drug differences in 9 out of the 11 subscales of the ASC (all but Anxiety and Impaired Control and Cognition) (**Supplementary Figure 8**). None of the subscales were significantly associated with redundancy PC. While the overall synergy and redundancy PC profiles were associated with phenomenological profiles across drug conditions, there were no significant correlations between specific network-level synergy or redundancy changes and specific phenomenological subscales, after multiple comparisons.

**Figure 5.**
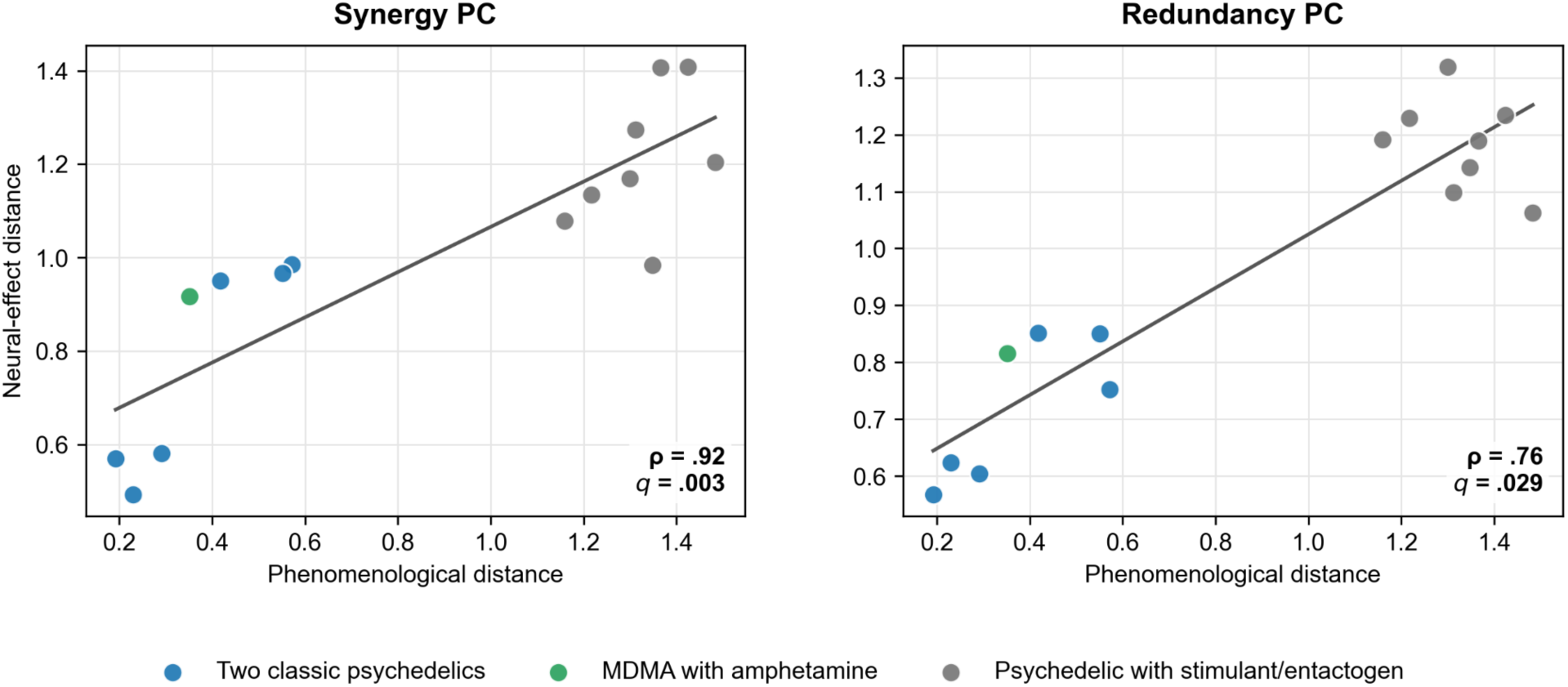
Between-drug differences in synergy and redundancy PC match between-drug differences in phenomenology. Phenomenology “profiles” were defined based on 5D-ASC ratings and correlated between drug conditions. Synergy PC and redundancy PC were also correlated between drug conditions, using prior methods (Figure 3). Both sets of correlations were converted into chord distances, which, unlike correlations, are Euclidean. Finally, between-drug phenomenological distances were compared with between-drug neural-effect distances for synergy PC and redundancy PC. The significance of this correlation was assessed with a Mantel test. The between-drug differences in synergy and redundancy PC were significantly similar to the between-drug differences in phenomenology.

## DISCUSSION

Here, we used an information-theoretic model to investigate how psychoactive drugs, including psychedelics, alter the architecture of the global workspace of consciousness. Classic psychedelics produced a distinct pattern across four drug conditions: they increased the number of gateways in the FPN and DAN. The spatial pattern of synergistic and redundant interactions distinguished classic psychedelics from stimulants/entactogens, while closely tracking between-drug differences in subjective experience. These findings suggest that classic psychedelics do not simply increase brain-wide integration. Instead, they reorganize how synergistic information is distributed across the global workspace.

Global workspace theory proposes that information becomes consciously accessible when it gains widespread access to distributed cortical systems.^36^ The global workspace has traditionally been described in terms of *ignition*: information that is initially processed within specialized systems crosses a threshold, becomes amplified through recurrent interactions, and gains access to higher-order cortex, which then distributes that information to the rest of the brain.^38–40^ This account has substantial support from the primate visual system.^51–55^ For example, van Vugt et al. (2018) subsequently showed that weak visual stimuli were consciously reported when activity survived successive stages of cortical processing and developed into strong, sustained frontal activity, a pattern that could be reproduced by a model of nonlinear ignition.^53^

The ΦID model, which was applied here, provides a complementary description of the synergistic global workspace.^42–44,56,57^ Rather than identifying ignition in response to a particular stimulus, it distinguishes workspace regions based on the ways in which they share information. Gateways are characterized by synergistic interactions that span a diverse set of networks, whereas broadcasters are characterized by broadly distributed redundant interactions. Across a series of studies, Luppi and colleagues have shown that the capacity for synergistic integration within this workspace is disrupted during loss of consciousness in humans and other mammals.^42–44^

Although related work has used similar information-theoretic techniques to quantify the effects of psychedelics on synergy,^58,59^ the present study, to our knowledge, is the first empirical application of the ΦID-derived synergistic global workspace to psychedelic states. Grinde & Stewart previously proposed a theoretical global-workspace account in which psychedelics loosen the selection or “gating” of information into conscious awareness, enabling novel or atypical content to enter the workspace.^60^ Our findings provide an empirical test of this proposal by showing that classic psychedelics reorganize two separable aspects of the global workspace: synergistic integration (i.e., gateways) and redundant dissemination (i.e., broadcasters). In particular, classic psychedelics were associated with increases in the number of gateways within networks involved in flexible control and attentional selection, i.e., the FPN and DAN.^61–65^ The FPN is composed of “flexible hubs” whose brain-wide connectivity changes more across tasks than that of other networks, allowing the network to adapt its interactions according to current goals and context.^64,65^ Importantly, the FPN was predominantly composed of broadcasters in our reference HCP data. Psychedelics therefore shifted a subset of frontoparietal regions from a configuration dominated by broadly distributed redundant interactions toward one in which synergistic interactions were more broadly distributed across networks. One possible interpretation is that the shift toward FPN gateways supports more flexible integration across brain systems.^44^ This would be consistent with evidence that psychedelics can increase neural flexibility and, more recently, that psilocybin enables brain dynamics to flexibly align with ongoing context, although psychedelic effects on behavioral flexibility vary across studies.^48,66–68^ Stoliker et al. (2026), who reported this context alignment, also found that psilocybin shifts integration toward associative and away from sensory regions rather than increasing it uniformly,^67^ which converges with our conclusion that classic psychedelics reorganize, rather than simply increase, integration. The accompanying increase in DAN gateways may complement the heightened neural flexibility on classic psychedelics because the DAN contributes to selecting and prioritizing information for attention.^61–63^ Increasing the synergistic integration within higher-order association networks may help to explain the unusual contents of states of consciousness induced by classic psychedelics. Such integration could condition sensory information, for example, more strongly on simultaneous emotional, attentional, or interoceptive states, which may give rise to synesthesia, atypical somatic experiences, hallucinations, and other abnormal subjective phenomena on classic psychedelics.^69,70^

Unlike classic psychedelics, MDMA and amphetamine did not both alter the number of gateways in higher-order networks (though MDMA alone did raise the number of gateways in the DAN). Instead, MDMA and amphetamine reduced the number of gateways within VIS and SMN, a pattern shared by some but not all classic psychedelics. This is consistent with prior ΦID work showing that redundant information is preferentially concentrated in unimodal sensorimotor cortex, whereas synergy is more prominent in higher-order association cortex.^44^ Both MDMA and amphetamine enhance norepinephrine signaling in part through substrate-mediated actions at the norepinephrine transporter, which increases extracellular norepinephrine.^71–73^ Norepinephrine neuromodulation can enhance signal-to-noise in sensory systems, including visual and somatosensory cortex, by suppressing background activity while preserving or enhancing stimulus-evoked responses.^74,75^ Such effects may make stimulus-related information more consistently represented across multiple regions, thereby increasing the amount of information that is available redundantly from different parts of the brain and favoring broadcaster-like processing; however, this link remains speculative. In the leave-one-network-out analysis, the effect of the SMN on redundancy PC, which is used to assign broadcasters, also distinguished classic psychedelics from MDMA and amphetamine. Unlike classic psychedelics, MDMA and amphetamine have prominent psychomotor stimulant effects and increase extroversion and verbal interaction: amphetamine increases motor activity, whereas MDMA increases psychomotor drive and can cause jaw clenching and restlessness.^76,77^ Thus, fewer SMN gateways on MDMA and amphetamine may reflect a broader distribution of the motor signals that are associated with the characteristic psychomotor effects of these drugs.

Overall, the spatial topography of synergy and redundancy PC distinguished not only between but also within drug classes. In particular, the DMN contributed to differences between LSD (LAM) and the LPM psychedelics in redundancy PC. Removing DMN regions significantly increased their similarity for redundancy PC; the corresponding increase for synergy PC did not survive FDR correction. Prior studies have claimed that multiple psychedelics all reduce the connectivity of the DMN,^16,18,78–81^ but our results provide a more nuanced computational account. Rather than producing a uniform DMN response, different classic psychedelics appear to distinctly reorganize how the DMN integrates information into the global workspace. For example, the number of DMN gateways increased under mescaline and LSD (LPM), but not under psilocybin or LSD (LAM); this pattern alone does not establish differences between drugs. Given the DMN’s involvement in autobiographical memory, internally generated cognition, and self-related processing, such variation could contribute to differences in the experiential or therapeutic effects of classic psychedelics.^82–85^

By contrast, the visual network showed a similar pattern across drug classes. For redundancy PC, removing VIS regions increased the separation of LSD from MDMA and amphetamine, but also reduced its similarity to the other classic psychedelics, indicating that the effects of the VIS made all these conditions more alike. Consistent with this result, both LSD conditions, MDMA, and amphetamine significantly reduced the number of gateways in VIS. This common effect may reflect a broad influence of psychoactive drugs on sensory processing, rather than a specific mechanism of psychedelic hallucinations. Indeed, MDMA and amphetamine do not produce the characteristic visual alterations of classic psychedelics,^14^ yet the two classes of drugs similarly affected the role of VIS in the global workspace. Hallucinations may instead arise when reductions in VIS gateways occur alongside gateway changes in higher-order networks.

Taken together, these findings show that classic psychedelics were distinguished from stimulants/entactogens by a distributed set of changes to the global workspace. Some features, such as the response of the VIS, were shared across drug classes; others, such as the contribution of the SMN, helped distinguish classic psychedelics from MDMA and amphetamine; and still others, particularly in the DMN, led to variation among classic psychedelics. The correspondence between the synergy/redundancy topographies and phenomenology profiles further suggests that the phenomenology of the drugs is associated with a distributed reconfiguration of the global workspace, rather than changes within any single network. Thus, the synergy and redundancy PCs are useful not only for classifying the drugs but also for characterizing the differences between the subjective experiences that they elicit. Drugs that produced similar perceptual changes also reorganized the global workspace in similar ways. Although previous studies have linked particular subjective experiences on psychedelics to greater connectivity between networks,^11,16,21,26,27,29,32,46–48^ here we show that the phenomenology of psychoactive drugs may instead depend on the spatial pattern of synergistic and redundant interactions in the global workspace. This provides a potential computational link between large-scale brain reorganization and the multidimensional structure of conscious experience, although it does not establish that these neural changes cause particular subjective effects.

Synergy PC corresponded to a larger number of 5D-ASC subscales than redundancy PC, potentially because synergistic interactions combine complementary information across systems supporting perception, emotion, cognition, and the self.^42,44^ (Note that this descriptive difference in significance counts does not establish stronger subscale associations for synergy PC than for redundancy PC.) The reorganization of synergistic interactions may relate to multiple interdependent aspects of psychedelic experience, whereas redundancy may reflect a less phenomenologically specific process of disseminating shared information. Nevertheless, the subscales were strongly correlated, and, across the six conditions, no subscale association remained significant after adjustment for the mean of the other ten subscales.

There are several limitations to acknowledge. Firstly, the workspace was defined from an independent HCP sample and held fixed across conditions, which assumes that workspace membership itself does not change under psychedelics. Second, between-drug comparisons may be influenced by cohort differences and by shared participants and placebo scans within each study. Although participant-level randomization tests addressed dependence within studies, replication across independent cohorts receiving the same drugs is needed, but only one mescaline fMRI dataset is currently available.^45^ Thirdly, in resting-state fMRI data, the ΦID model is unable to identify *which* information reaches the global workspace of consciousness. Therefore, the model cannot distinguish whether increases in the number of gateways specifically encode synesthesia, reliving of traumatic memories, ego dissolution, or any of the other myriad subjective experiences that can occur on psychedelics. Fourth, differences in synergy and redundancy between brain states can be confounded by differences in total mutual information: in MEG data from LSD, psilocybin, and ketamine, raw synergy was lower in the drug state, whereas synergy normalized against null models was higher and normalized redundancy lower.^59^ Our main outcomes are less exposed to this confound than raw synergy and redundancy. Participation coefficients of synergy and redundancy, which are used to define gateways and broadcasters, depend on how each region’s synergistic or redundant interaction strength is distributed across networks rather than on its overall magnitude.

## CONCLUSION

Previous studies have demonstrated that psychedelics increase global connectivity in the brain,^15,86,87^ and recent work indicates that psilocybin redistributes integration from sensory toward associative cortex.^67^ Our work extends these findings to the informational architecture of the global workspace of consciousness. We showed that psychedelics do not simply increase the integration of information within the global workspace. Instead, classic psychedelics increased the number of gateways in specific higher-order networks, i.e., the FPN and DAN, while broadening the distribution of both synergistic and redundant interactions across brain networks. Moreover, the regional topographies of synergy and redundancy PC distinguished classic psychedelics from stimulants/entactogens, while differences in these topographies across drugs corresponded to differences in phenomenology. Together, these findings provide a computational account of how the architecture of the global workspace may contribute to the unique character of the psychedelic experience.

## METHODS

### Datasets and participants

We performed secondary analyses of three resting-state fMRI datasets. The Human Connectome Project 100-unrelated subsample (*n* = 100 healthy adults, one resting-state run each) was used to define the global workspace.^49^ The LAM study, conducted at the University of Basel, administered LSD, D-amphetamine, MDMA and placebo in a within-subjects crossover design (each participant received each drug).^14^ The LPM study, conducted at the University of Basel, administered LSD, psilocybin, mescaline and placebo, also in a within-subjects crossover design.^45^ The two Basel studies were registered three years apart and share no participants.

After motion quality control, we analyzed data from *n* = 26 participants (LSD, amphetamine, and MDMA) in the LAM study and *n* = 30 (LSD), *n* = 31 (psilocybin), and *n* = 31 (mescaline) participants in the LPM study. For each condition, an equivalent number of participants were analyzed in the placebo control. All participants in all three studies were healthy. Characteristics of participants, dosing and design are in **Table 1**. Note that the mescaline arm pools two doses, since half of the participants received 300 mg while the other half received 500 mg in the parent study.^45^

**Table 1.**
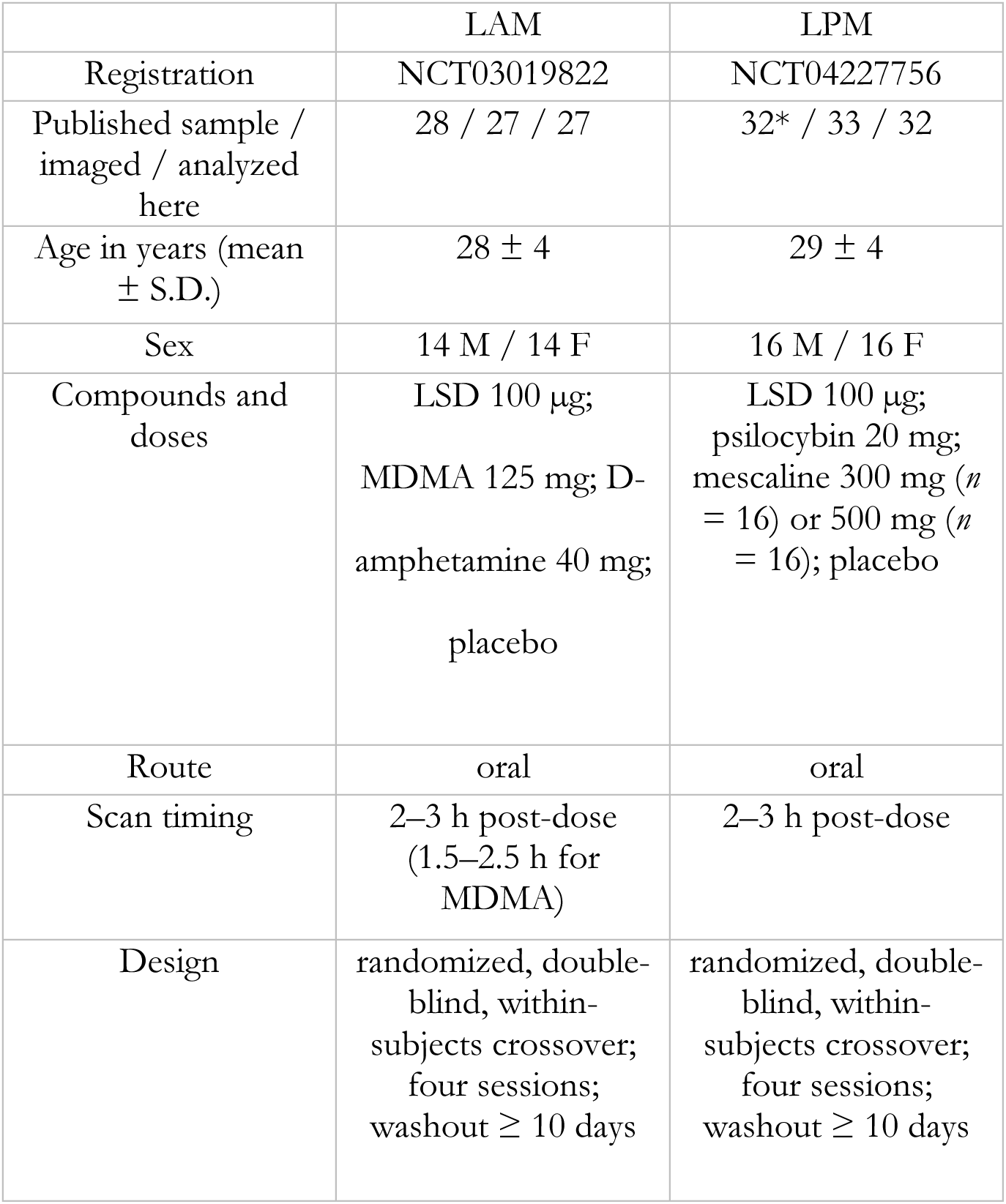
Characteristics of participants and study design. Note that “analyzed here” is the number of participants contributing at least one usable drug-control pair of scans. The Methods give the per-condition *n* after motion screening, which is lower. *The parent study had 32 participants who completed the trial, but 33 total people were imaged.

Both studies were conducted in accordance with the Declaration of Helsinki and approved by the Ethics Committee of Northwest Switzerland. The Swiss Federal Office for Public Health authorized the administration of LSD, amphetamine, MDMA, psilocybin, and mescaline to healthy participants.

### Image acquisition and preprocessing

Acquisition parameters from each dataset are described in detail in **Supplementary Table 5**. The LAM and LPM studies share the same exact scanner and acquisition parameters. All fMRI data from all three studies were preprocessed with an identical pipeline. Preprocessing used fMRIPrep 23.2.1.^88,89^ BOLD images underwent head-motion correction, coregistration to the participant’s T1-weighted anatomical image, and normalization to MNI152NLin2009cAsym space. Slice-timing correction was performed using AFNI 3dTShift when valid slice-timing metadata were available and susceptibility-distortion correction was applied when supported by suitable field-map acquisitions and associated metadata. No spatial smoothing was applied. After removal of dataset-specific initial volumes, we excluded scans exceeding 3 mm maximum absolute translation or 3.15° maximum absolute rotation, truncating four scans to a clean segment rather than dropping them.^42^ Drug effects on network gateway counts and mean synergy and redundancy PCs were re-estimated with each participant’s drug-minus-control change in framewise displacement partialed out. For the motion-adjusted analyses, mean framewise displacement was calculated over the retained volumes used for ΦID, including the retained segment of each truncated scan.

Images were then denoised using CONN v25.b.^90^ CONN segmented each participant’s preprocessed T1 image to define white-matter and cerebrospinal-fluid compartments. Nuisance regressors comprised five anatomical CompCor components from each compartment, six fMRIPrep realignment parameters, and the six corresponding first derivatives.^91^ Linear detrending and 0.008–0.09 Hz temporal filtering were applied. No despiking, isolated-frame censoring, ART scrubbing, or global signal regression was performed.

Denoised BOLD signals were spatially averaged within 454 regions comprising the Schaefer-400 cortical and Tian-54 subcortical atlases in MNI152NLin2009cAsym space.^92,93^ The Schaefer-400 parcellation contains seven networks: the visual network (VIS), somatomotor network (SMN), dorsal attention network (DAN), ventral attention network (VAN), limbic network (LIM), frontoparietal network (FPN), and default mode network (DMN).^94^ Parcel time series were deconvolved to estimate latent neural signals using the rsHRF toolbox v2.5 prior to ΦID, thereby reducing the influence of regional hemodynamic response differences on time-delayed information estimates.^42,95^

### Integrated information decomposition

For every pair of the 454 regions, the time-delayed mutual information between the pair’s past and future (lag = 1 TR) was decomposed with integrated information decomposition under the minimum-mutual-information redundancy function for Gaussian variables.^42,43^ The decomposition splits the information that two regions’ present carries about their joint future into components: information available redundantly from either region alone, and information available only synergistically, from both together. From this decomposition, we used the persistent-synergy atom (i.e., past synergy to future synergy) to define synergy between pairs of regions and the persistent-redundancy atom (i.e., past redundancy to future redundancy) to define redundancy between pairs of regions.

### Global workspace, gateways, and broadcasters

Following the methods of Luppi et al. (2024), the global workspace was defined once, based on the 100 HCP participants.^42^ For each region, node strength was computed on the group-average synergy and redundancy matrices; regions whose synergy-strength rank exceeded their redundancy-strength rank constitute the global workspace. There are 235 regions in the global workspace. Within the workspace, regions were classified by PC with respect to eight modules: the seven cortical networks of the parcellation plus one subcortical module.^96^ Gateways are defined as regions within the global workspace in which the synergy PC rank exceeds the redundancy participation coefficient rank; broadcasters are regions within the global workspace in which the redundancy participation coefficient rank exceeds the synergy PC rank.

### Drug-induced changes in gateway counts per network

Using two-sided paired t-tests, we tested within-subject drug-versus-control differences in the number of regions classified as gateways within each resting-state network. Regions with equal synergy and redundancy PC ranks are assigned to neither role, so gateway and broadcaster counts need not change by exactly equal amounts. Two families of tests were defined a priori. Gateway-count changes in the FPN and DMN constituted the primary family, as motivated in the Introduction, while changes in the remaining six networks constituted the exploratory family. Within each drug condition, FDR correction was applied separately across the two primary tests and across the six exploratory tests. Correction was not performed across drug conditions.

### Drug-induced changes in synergy and redundancy PCs

Whereas the preceding analysis measured changes in gateway counts within networks, we also examined changes in mean synergy and redundancy PC. Each subject’s synergy and redundancy PCs were averaged across the 235 workspace regions, and drug-versus-control differences were tested with two-sided one-sample t tests on the paired differences. FDR correction was applied across the six drug conditions separately for synergy PC and redundancy PC; correction was not applied across the two measures. Effect sizes were expressed as bias-corrected paired-samples Hedges’ g_z_.

### Separating classes of drugs

The six drug conditions belong to two pharmacological classes: classic psychedelics (LSD, psilocybin, and mescaline) and stimulants/entactogens (amphetamine and MDMA). For each drug condition, drug-versus-placebo effects on synergy PC and redundancy PC were represented by 235-region vectors of Hedges’ *g*_av_ values. Hedges’ *g*_av_ was used because these vectors quantify standardized differences between repeated-measures conditions while retaining the variability of each condition. Pearson correlations between these regional effect profiles quantified the similarity between drug conditions.

Classification was evaluated using one-sided participant-level randomization tests in the hypothesized directions. These analyses used participants with usable data from all four sessions (LAM: *n* = 24; LPM: *n* = 28). Within each study, drug condition labels were randomly permuted separately for every participant, while the placebo condition was held fixed. Regional effect profiles for synergy and redundancy PC and the corresponding test statistics were recomputed for 100,000 randomizations. Three complementary sets of tests were performed. First, within LAM, separation was defined as the MDMA-amphetamine similarity minus the mean similarity of LSD with MDMA and amphetamine; larger values indicate stronger separation of LSD from the stimulant/entactogen conditions. Second, cross-study psychedelic similarity was defined as the mean similarity of LSD from LAM with LSD, psilocybin, and mescaline from LPM; larger values indicate greater-than-expected similarity across studies. Third, cross-study class separation tested whether the mean similarity between the LPM psychedelics and MDMA or amphetamine was lower than expected.

For reporting and display, the permutation-null mean was subtracted from the observed statistic in the first two sets; in the third set, the reported statistic was the null mean minus the observed similarity. Thus, zero represents the expectation under randomized condition labels. Positive values indicate greater class separation in the first and third sets of tests and greater cross-study psychedelic similarity in the second set of tests. FDR correction was applied across the two within-LAM tests (synergy PC and redundancy PC) and, separately, across the four cross-study tests comprising both PC measures in the second and third sets.

For descriptive visualization, drug conditions were also clustered according to the similarity of their synergy and redundancy PCs. Within-study condition profiles were embedded in two dimensions using classical multidimensional scaling of the chord distances derived from their Pearson correlations. The resulting layouts were used only to illustrate the relationships quantified by the randomization tests and were not treated as inferential clustering analyses. Accordingly, the shaded groupings indicate the pharmacological classes being tested rather than statistically supported clusters. Line thickness represents profile similarity on a common nonlinear scale across panels, with thicker lines indicating stronger positive correlations.

To determine which networks contributed to drug-class separation, we repeated each classification analysis after removing, in turn, all workspace regions belonging to each of the eight networks. For each omitted network, its entries were removed from the regional effect profiles, and the relevant similarity or separation statistic was recalculated; the original participation coefficients were held fixed. Participant-level bootstrap resampling was used to derive 95% confidence intervals for these changes.

### Comparing global workspace to phenomenology

Finally, we examined whether differences among drugs in the regional topographies of synergy PC and redundancy PC corresponded to differences in subjective experience. Phenomenological profiles comprised mean scores on the 11 lower-order 5D-ASC scales described by Studerus et al.; the questionnaire was administered in both the LAM and LPM studies (**Supplementary Table 6**).^50^ The analysis included six drug conditions: LSD (LAM), LSD (LPM), psilocybin, mescaline, MDMA, and amphetamine.

For each neural measure, every drug condition was represented by a 235-region vector of bias-corrected effects relative to its corresponding placebo condition, standardized using the pooled within-condition standard deviation. Neural profiles used available drug–placebo pairs: *n* = 26 per LAM contrast and *n* = 30, 31, and 31 for LSD, psilocybin, and mescaline, respectively, in LPM. Similarity between each pair of conditions was quantified using Pearson correlation across the 235 regional effects. Phenomenological similarity was likewise calculated using Pearson correlation across the eleven mean 5D-ASC subscale scores. Similarities were converted to chord distances as *d_ij_* = 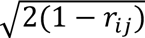 where *rᵢⱼ* is the Pearson correlation between conditions *i* and *j*. This placed the neural and phenomenological relationships on a common Euclidean distance scale, with larger values indicating greater dissimilarity. Separate 6 x 6 distance matrices were produced for phenomenology, synergy PC, and redundancy PC, each containing 15 unique off-diagonal distances.

Mantel tests assessed whether pairs of conditions that were more phenomenologically dissimilar were also more dissimilar in their regional neural-effect profiles. Correspondence was quantified using Spearman’s ρ across the 15 matched off-diagonal distances. Statistical significance was evaluated by exhaustively permuting the six condition labels of the phenomenological matrix while preserving its internal pairwise structure, yielding all 6! = 720 possible relabelings. Two-sided exact *p*-values were calculated as the proportion of permutations producing an absolute correlation at least as large as the observed value. FDR correction was applied across the two PC measures.

As a follow-up analysis, we repeated the Mantel test separately for each of the eleven ASC subscales to identify which dimensions of subjective experience contributed to the overall correspondence with neural geometry. FDR correction was applied separately across the eleven subscale tests for synergy PC and across the eleven tests for redundancy PC. As an exploratory sensitivity analysis, each subscale’s six condition means were regressed on an intercept and the mean of the other ten subscales. Mantel tests were repeated using pairwise absolute differences between the residuals, with FDR correction across eleven subscales separately for each PC measure and separately from the unadjusted tests.

### Use of artificial intelligence

OpenAI ChatGPT and Anthropic Claude were used to assist with writing and debugging analysis code. All AI-assisted code was reviewed and tested by the authors, who take responsibility for the final code and analyses. Additionally, ChatGPT and Claude were used to review and edit the text of the manuscript.

## ACKNOWLEDGMENTS

K.S. is funded by a War Related Illness and Injury Study Center Fellowship from the Veterans Affairs Palo Alto Healthcare System. The LAM and LPM studies, for which data was analyzed in this study, were supported by the Swiss National Science Foundation (grant no. 320030_170249 and grant no. 32003B_185111 to M.E.L., respectively). The LPM study was additionally supported by the University Hospital Basel and Mind Medicine Inc.

## AUTHOR CONTRIBUTIONS

P.Y. preprocessed and analyzed the data and assisted with writing the manuscript. K.S. conceptualized the project, assisted with data analysis, and wrote the manuscript. P.V., F.H., A.M.B., and F.M. acquired the data. M.E.L. designed the LAM and LPM studies. G.H., C.I.R., A.L., and P.A.M.M. edited the manuscript.

## DATA AND CODE AVAILABILITY

Data may be available upon request by contacting F.M. All code for this study is available online: doi.org/10.5281/zenodo.22677151.

## DECLARATION OF COMPETING INTERESTS

M.E.L. was a consultant for Definium Therapeutics and Resilient Therapeutics. F.M. was a consultant for Definium Therapeutics. Know-how and data associated with this work are owned by the University Hospital Basel and were licensed by Definium Therapeutics. Definium Therapeutics had no role in the design, conduct, analysis, interpretation of the data, or the decision to publish the present study.

## SUPPLEMENTARY FIGURES AND TABLES

**Supplementary Figure 1.**
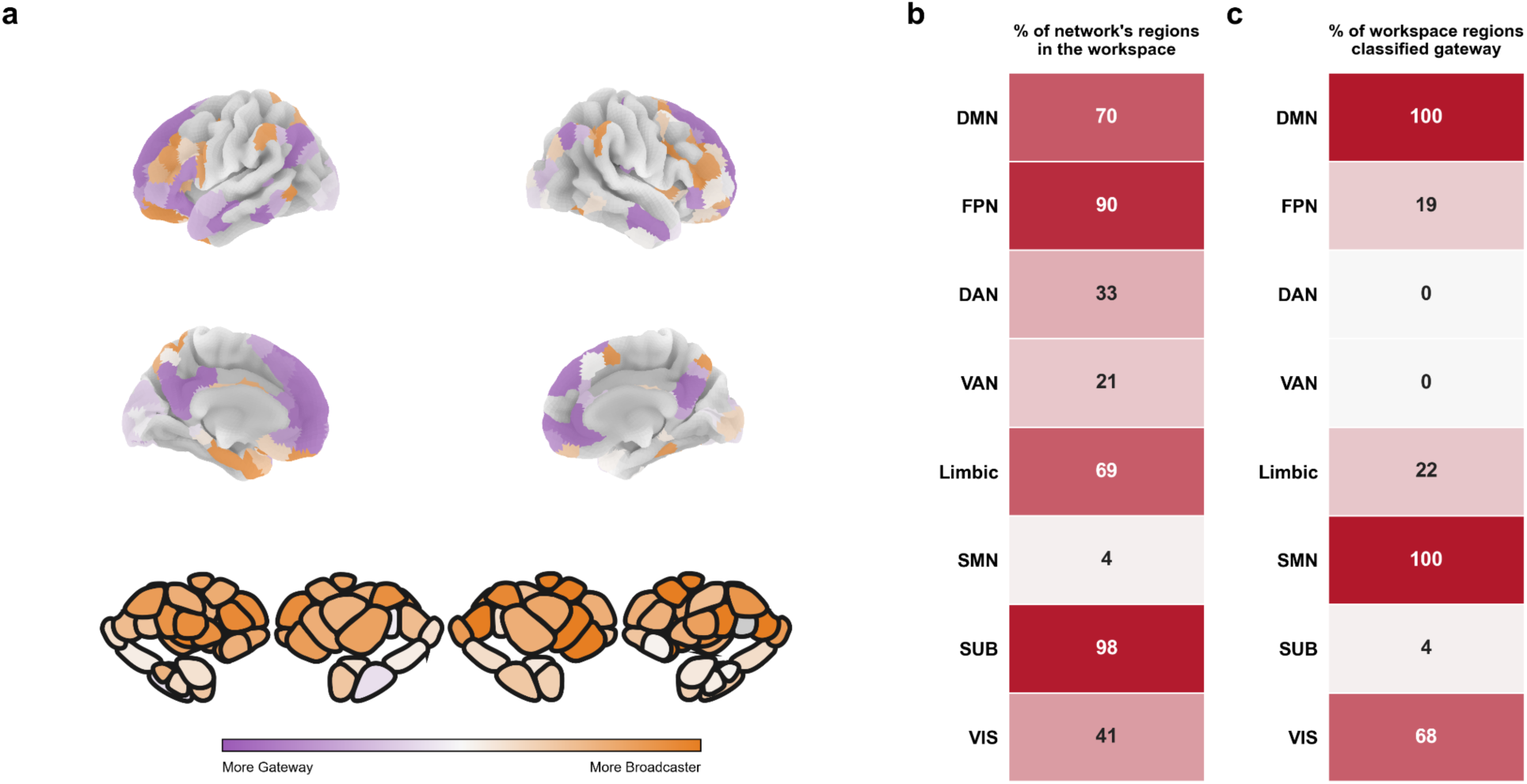
The global workspace, defined on the HCP data. (a) Gateway (violet) and broadcaster (orange) assignments of the 235 workspace regions in the HCP sample (*n* = 100); grey regions are outside the workspace. Beneath the four views of the cortex, the subcortex is shown. The more violet a region, the higher its within-workspace synergy PC rank relative to its redundancy PC rank; the more orange a region, the higher its redundancy PC rank relative to its synergy PC rank. (b) Percentage of each network’s regions included in the workspace. (c) Percentage of each network’s workspace regions classified as gateways. All eight modules of the participation partition are shown (seven Yeo cortical networks plus subcortex).

**Supplementary Figure 2.**
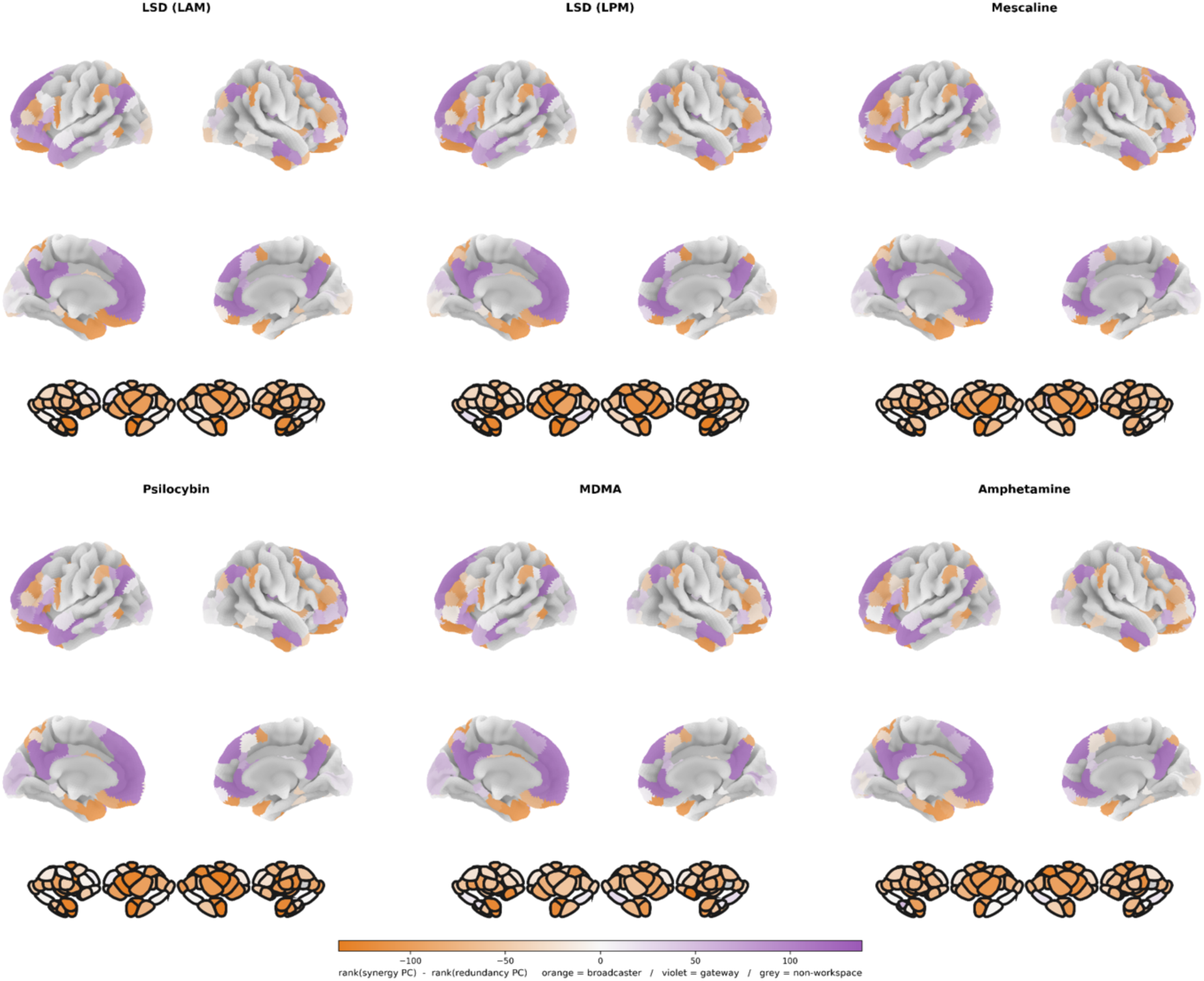
Topography of gateways and broadcasters in each of the six drug conditions. Within the 235-region HCP-defined workspace, color represents rank(synergy PC) − rank(redundancy PC): positive differences indicate gateways (violet), negative differences indicate broadcasters (orange), and equal ranks are assigned to neither role. Grey regions are outside the workspace. For each condition, four views of the fsaverage5 pial cortical surfaces and a Tian S4 subcortical strip are shown. A common color scale is used across the six conditions. The topography of gateways and broadcasters is largely consistent under each drug condition, but significant changes in the distribution of gateways are shown in **Figure 2**.

**Supplementary Figure 3.**
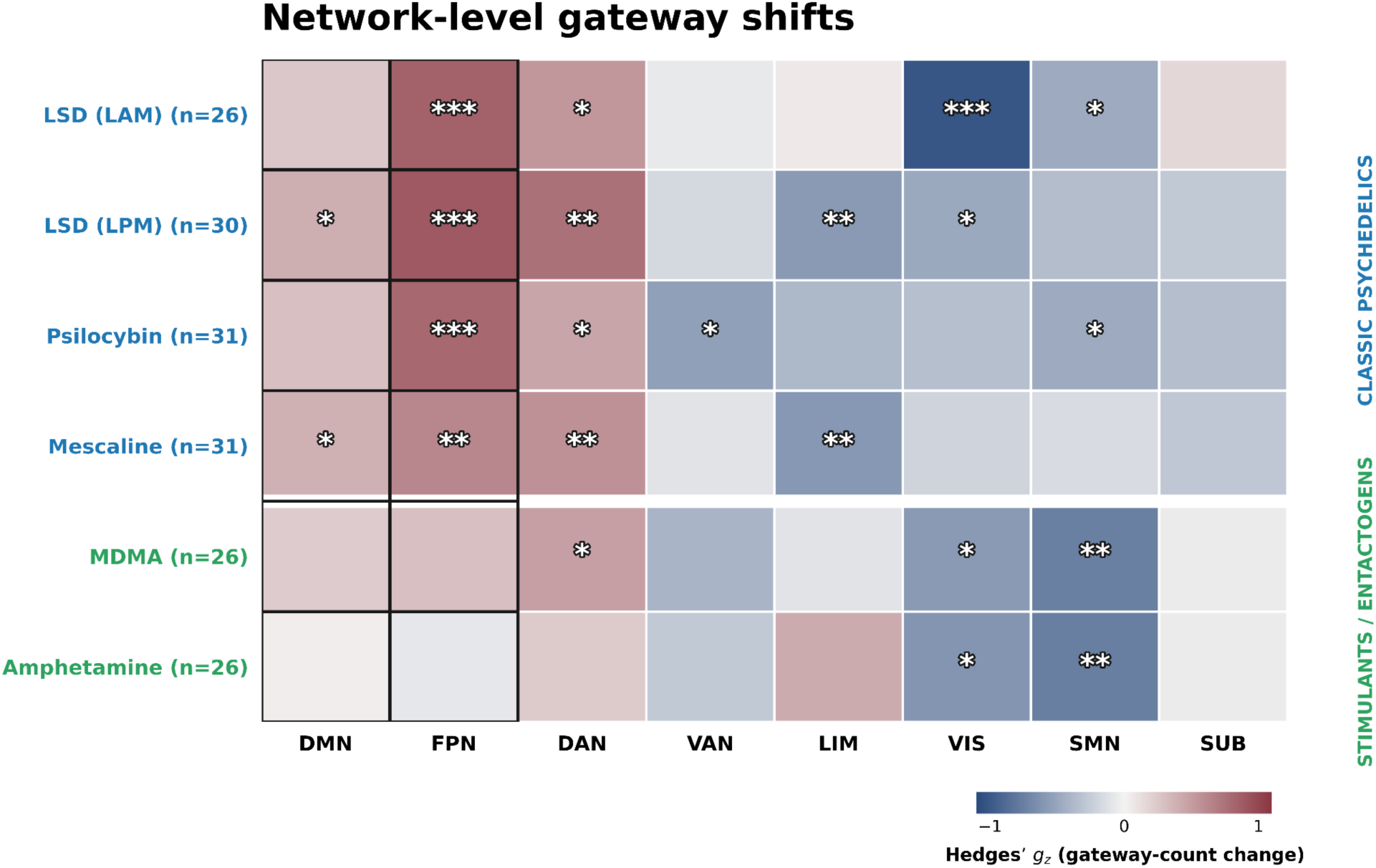
Effect of psychoactive drugs on gateway counts across networks. Each cell shows the standardized within-subject drug-versus-control change in gateway count for one resting-state network, expressed as bias-corrected paired-samples Hedges’ *g*_z_. Positive values indicate increases in the number of gateways in a given network, whereas negative values indicate decreases in the number of gateways in a given network. Gateway counts were tested once per network; broadcaster counts were not tested separately because the two outcomes are nearly reciprocal. Asterisks indicate significant two-sided paired *t*-tests after FDR correction. Correction was performed separately within each drug condition across the two primary networks, DMN and FPN, and across the remaining six exploratory networks; correction was not performed across drug conditions. Outlined cells indicate the primary networks. All four classic-psychedelic conditions increased FPN and DAN gateway counts, while LSD (LPM) and mescaline also increased DMN gateway counts. MDMA and amphetamine both reduced gateway counts in the VIS and SMN networks. * *q* < .05, ** *q* < .01, and *** *q* < .001.

**Supplementary Figure 4.**
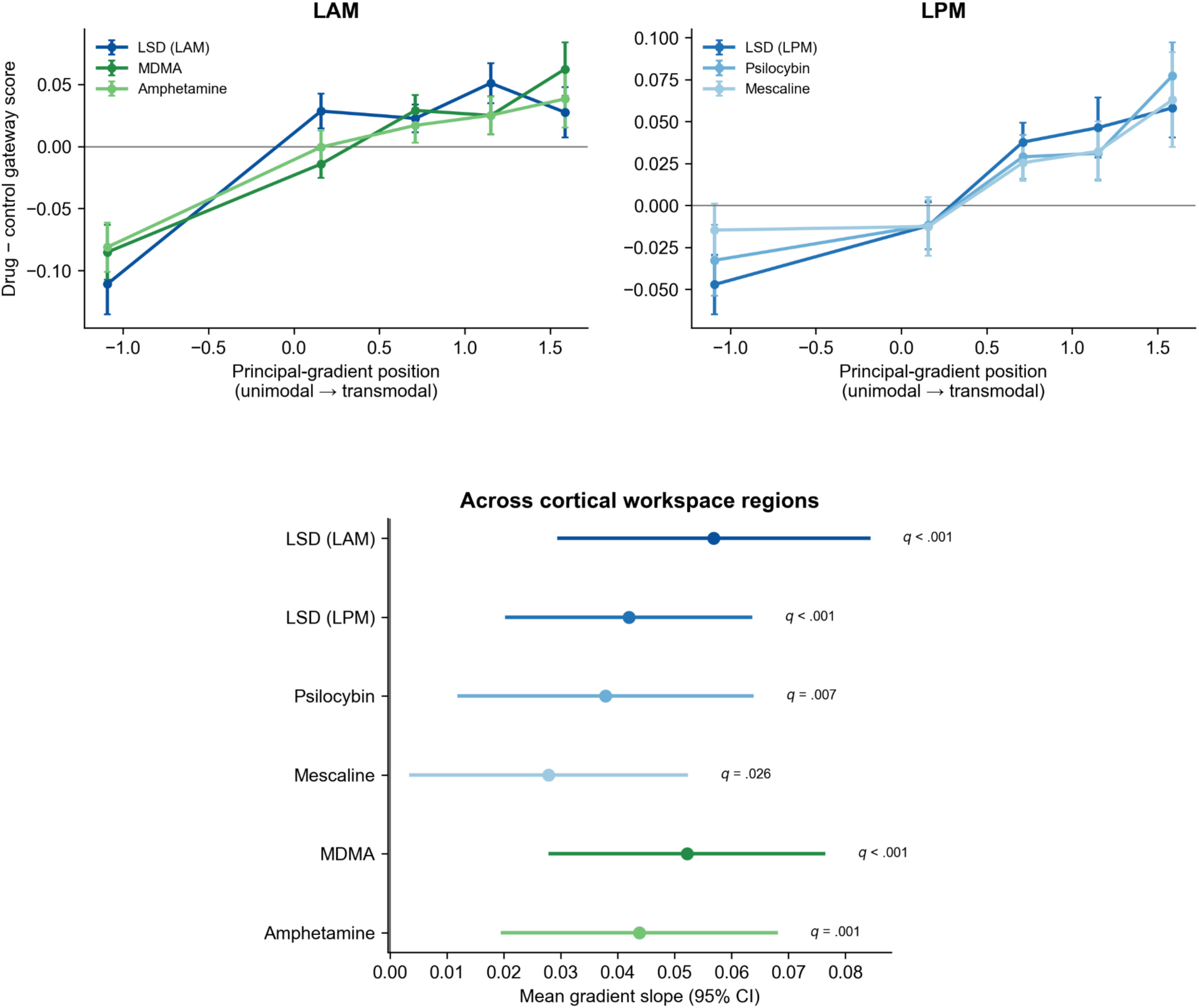
Psychoactive drugs tend to increase gateways in higher-order brain regions. Here, the gateway “score” is the normalized difference between each region’s within-workspace rank in synergy PC and redundancy PC. The principal connectivity gradient was derived independently from group-average Schaefer-400 functional connectivity. Lower-order regions on this gradient are in unimodal cortex (e.g., visual and somatomotor networks), whereas higher-order regions on this gradient are in transmodal cortex (e.g., frontoparietal control network and default mode network). Top panels show mean drug-versus-control changes in gateway score across five bins containing approximately equal numbers of cortical workspace regions. Error bars indicate the standard error across participants. Bottom panel: points show the mean participant-level slope relating gateway score changes to gradient position, with 95% confidence intervals. Positive slopes indicate greater drug-related increases in gateway scores in higher-order regions. Significance was assessed using two-sided participant-level sign-flipping tests, with FDR correction applied across the six drug conditions separately for each analysis.

**Supplementary Figure 5.**
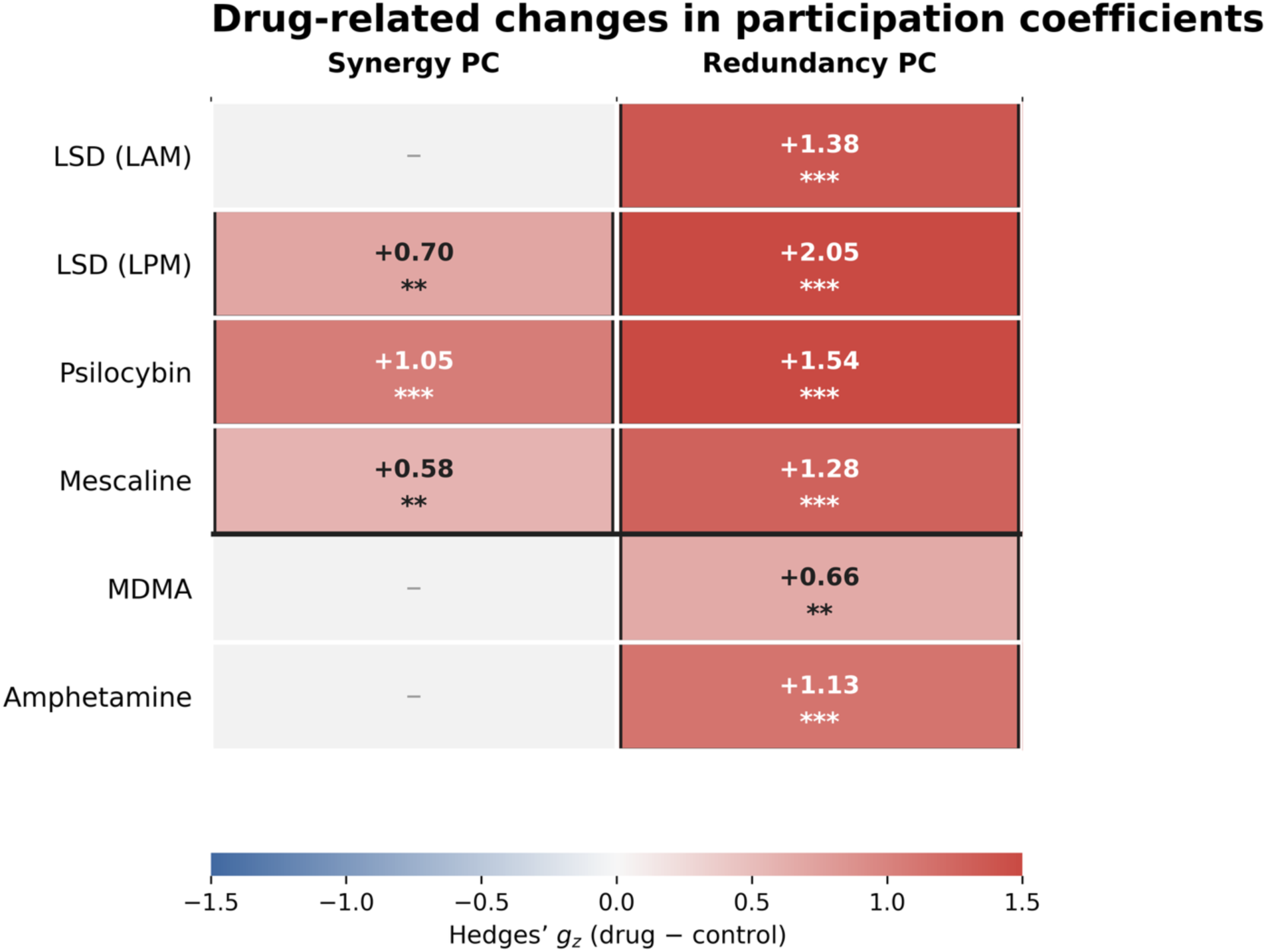
Drug-related changes in mean synergy and redundancy PCs. Synergy and redundancy PCs were averaged across the 235-region global workspace and compared between each active condition and its corresponding control condition using two-sided paired t-tests. Effects are expressed as bias-corrected paired-samples Hedges’ *g*_z_, with positive values indicating higher PCs under the active condition. FDR correction was applied across the six drug conditions separately within each PC measure; correction was not applied across the two measures. Colored cells show significant effects after correction, whereas dashes indicate nonsignificant effects. Redundancy PC increased significantly in all drug conditions. Synergy PC increased significantly under LSD (LPM), psilocybin, and mescaline, but not under LSD (LAM), MDMA, or amphetamine. * *q* < .05, ** *q* < .01, and *** *q* < .001.

**Supplementary Figure 6.**
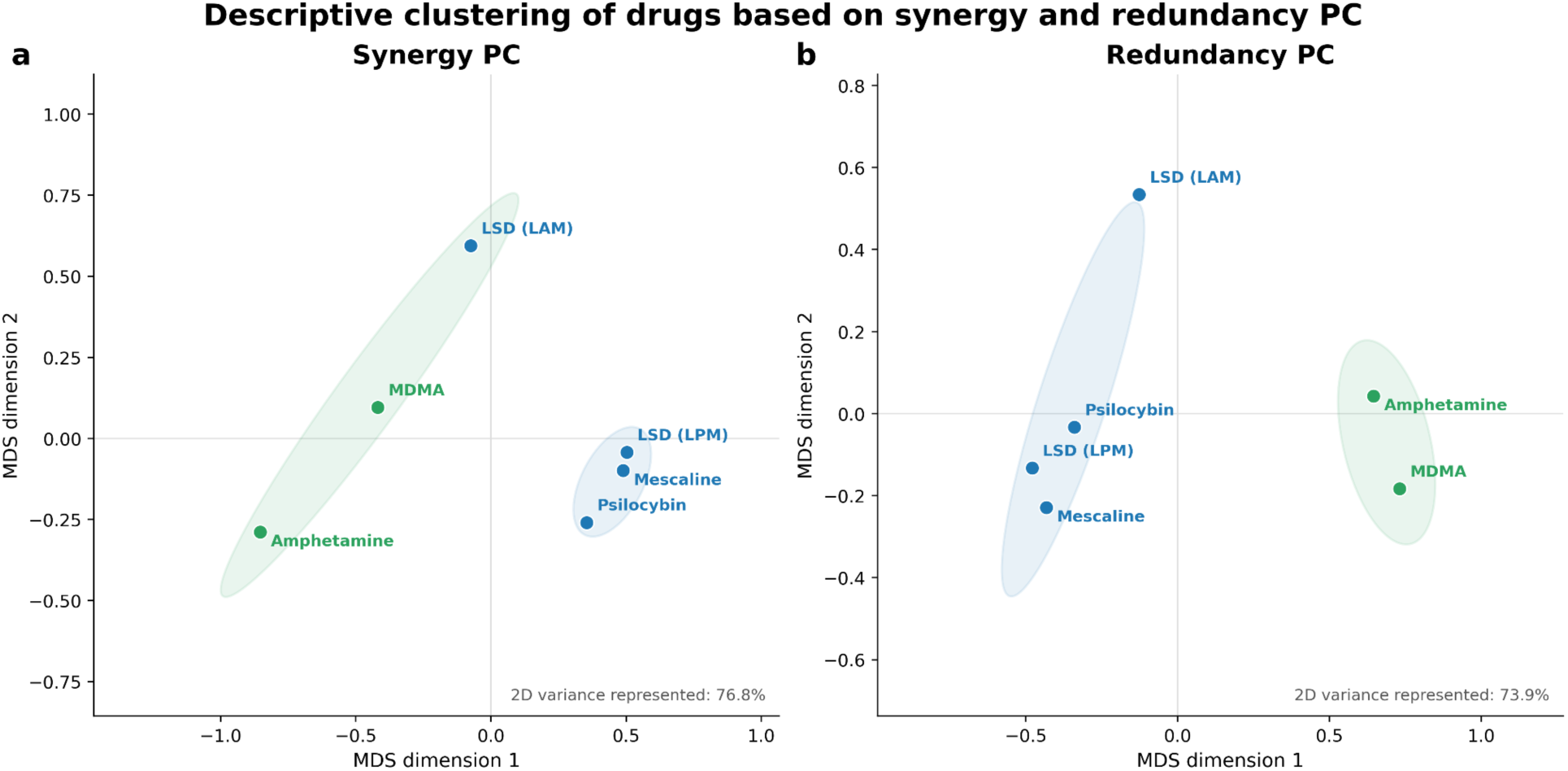
Descriptive clustering of drugs based on synergy and redundancy PC. The difference in synergy and redundancy PC between each drug condition and placebo was expressed as paired-samples Hedges’ *g*_av_ values. Pearson correlations between profiles were converted to chord distances. Ward’s minimum-variance hierarchical clustering used the full chord-distance geometry to define two clusters, and classical multidimensional scaling (MDS) provided the two-dimensional visualization. The two displayed dimensions represented 76.8% of the synergy-PC geometry and 73.9% of the redundancy-PC geometry. Shaded ellipses indicate the two-cluster Ward solution rather than confidence regions. For synergy PC, the three LPM psychedelics clustered together, whereas LSD (LAM) clustered with MDMA and amphetamine; this two-cluster solution was recovered in 99.3% of 10,000 participant-bootstrap resamples. For redundancy PC, all four classic-psychedelic conditions clustered separately from MDMA and amphetamine, with only 59.1% participant-bootstrap support. These clusters are descriptive because conditions acquired within the same study shared participants and placebo scans.

**Supplementary Figure 7.**
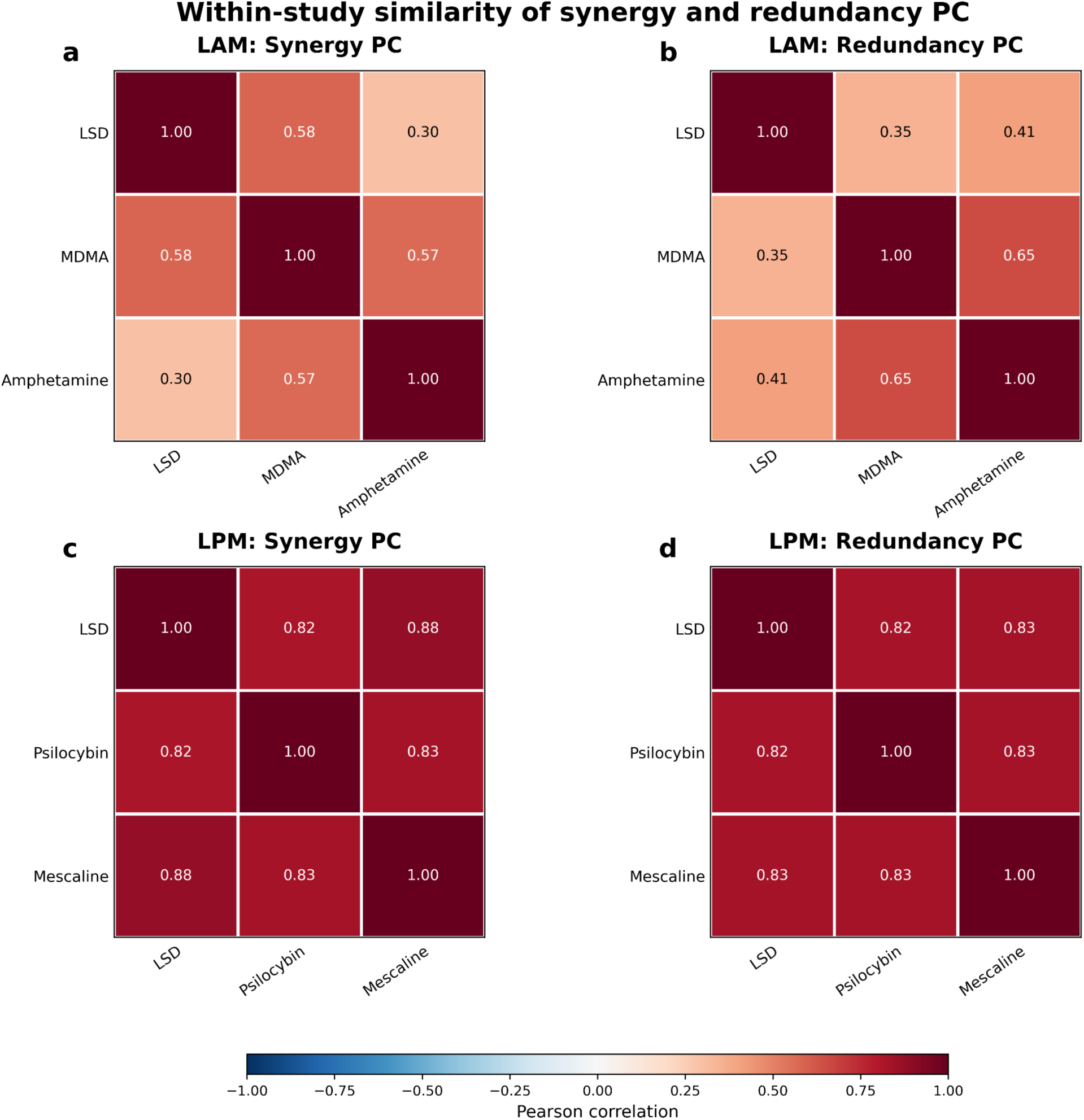
Between-drug correlations in synergy and redundancy PC. The difference in synergy and redundancy PC between each drug condition and placebo was expressed as paired-samples Hedges’ *g*_av_ values. These values were then correlated between drugs in each study. Each off-diagonal entry in the figure is one of these correlations. Synergy and redundancy PC are highly correlated between classic psychedelics, but not between classic psychedelics (e.g., LSD) and other drugs (e.g., amphetamine).

**Supplementary Figure 8.**
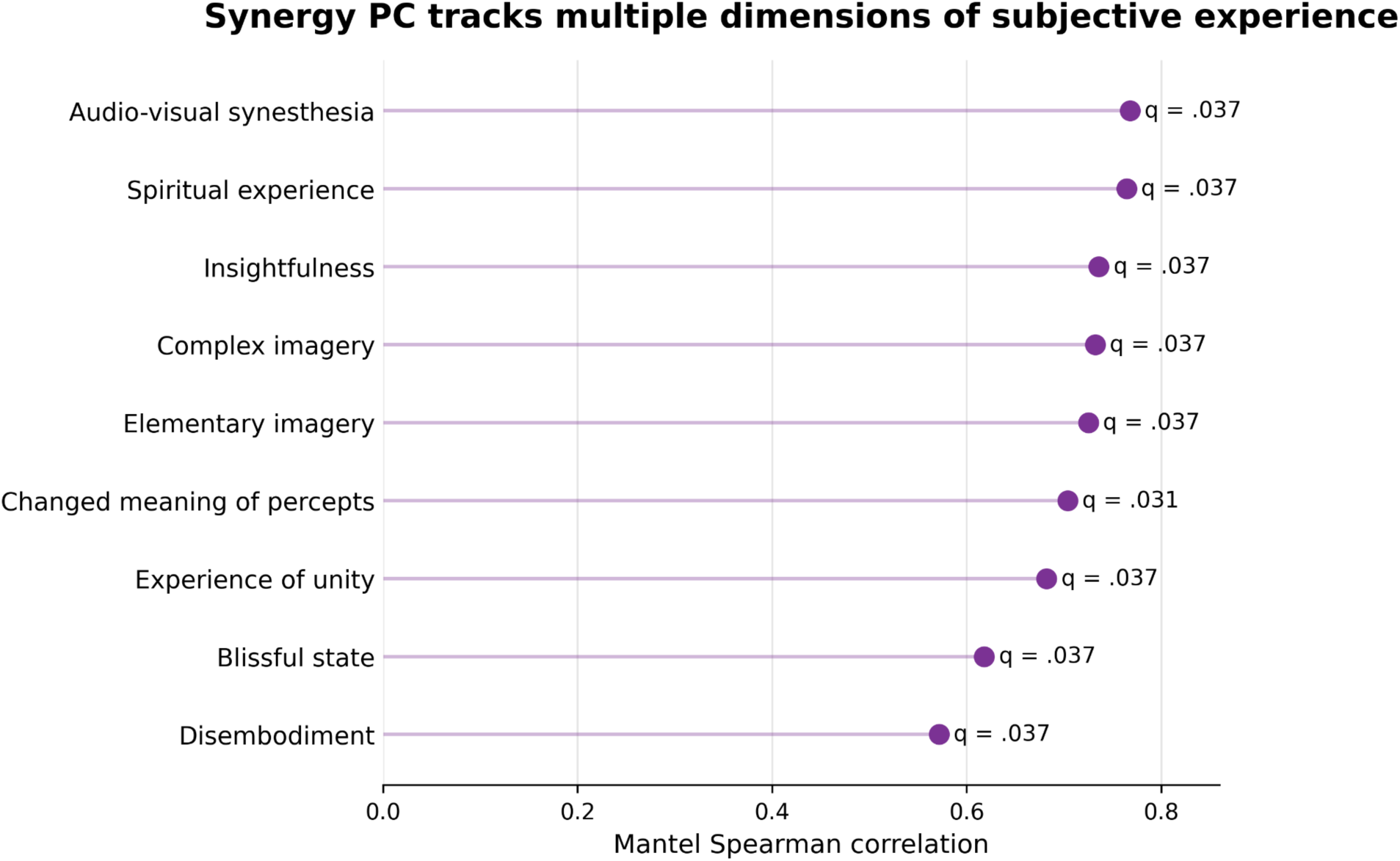
Between-drug differences in synergy PC correspond to multiple dimensions of subjective experience. Synergy PC and redundancy PC were correlated between drugs, and these correlations were converted to chord distances. For each 5D-ASC subscale, phenomenological distance was defined as the absolute difference in the mean subscale score between each pair of conditions. Mantel Spearman correlations tested whether condition pairs with more dissimilar synergy-PC effects also differed more strongly in each subjective dimension. Significance was assessed relative to all 720 condition-label permutations, with FDR correction across the 11 subscales. Only associations surviving this correction are shown. Between-drug differences in redundancy PC were not significantly associated with between-drug differences in any of the 5D-ASC subscales, whereas between-drug differences in synergy PC were significantly associated with between-drug differences in most of the 5D-ASC subscales.

**Supplementary Table 1.** Number of gateways per brain network in each of the drug and placebo conditions. For each condition, synergy and redundancy matrices were averaged across available retained scans before computing participation coefficients and assigning gateway roles within the fixed HCP-defined workspace. Abbreviations: VIS = visual network; SMN = somatomotor network; DAN = dorsal attention network; VAN = ventral attention network; LIM = limbic network; FPN = frontoparietal control network; DMN = default mode network; SUB = subcortical; HCP = Human Connectome Project; AMPH = D-amphetamine; PSI = psilocybin; MES = mescaline.

|  | VIS | SMN | DAN | VAN | LIM | FPN | DMN | SUB | Total gateways |
| --- | --- | --- | --- | --- | --- | --- | --- | --- | --- |
| HCP | 17 | 3 | 0 | 0 | 4 | 9 | 64 | 2 | 99 |
| LAM placebo | 22 | 3 | 0 | 0 | 3 | 13 | 61 | 0 | 102 |
| LPM placebo | 14 | 3 | 0 | 0 | 3 | 7 | 61 | 1 | 89 |
| LSD (LAM) | 12 | 3 | 0 | 0 | 2 | 23 | 63 | 3 | 106 |
| AMPH | 13 | 3 | 0 | 0 | 3 | 8 | 61 | 2 | 90 |
| MDMA | 17 | 3 | 0 | 0 | 3 | 10 | 62 | 4 | 99 |
| LSD (LPM) | 8 | 3 | 0 | 0 | 3 | 20 | 64 | 2 | 100 |
| PSI | 17 | 3 | 0 | 0 | 2 | 16 | 63 | 1 | 102 |
| MES | 15 | 3 | 0 | 0 | 3 | 16 | 62 | 1 | 100 |

**Supplementary Table 2.** Changes in the number of gateways per network for each drug condition relative to placebo/control. ‘M control’ and ‘M drug’ are the means of participant-level gateway counts in the matched control and drug samples, respectively. Δ is M drug − M control. Regions with equal synergy and redundancy PC ranks are assigned to neither role, so gateway and broadcaster counts need not change by exactly equal amounts. Drug-versus-control differences were tested using two-sided paired *t*-tests. Bias-corrected paired-samples Hedges’ *g*_z_ values and 95% confidence intervals are reported as effect sizes. FPN and DMN constituted the prespecified primary family (P), while the remaining six networks constituted the exploratory family (E). Within each drug condition, FDR correction was applied separately across the two primary tests and the six exploratory tests; correction was not performed across drug conditions. Abbreviations: VIS = visual network; SMN = somatomotor network; DAN = dorsal attention network; VAN = ventral attention network; LIM = limbic network; FPN = frontoparietal control network; DMN = default mode network; SUB = subcortical; AMPH = D-amphetamine; PSI = psilocybin; MES = mescaline.

| Drug cond. | Network | M control | M drug | $\Delta$ | $t$ | df | $p$ | Hedges' $g$,<br>[95% CI] | Tier | $q$ (tier) | FDR sig. |
| --- | --- | --- | --- | --- | --- | --- | --- | --- | --- | --- | --- |
| LSD (LAM) | DMN | 54.81 | 55.69 | 0.88 | 1.33 | 25 | .196 | 0.25<br>[-0.13, 0.63] | P | .196 | No |
| LSD (LAM) | FPN | 15.77 | 20.12 | 4.35 | 4.41 | 25 | < .001 | 0.84<br>[0.39, 1.27] | P | < .001 | Yes |
| LSD (LAM) | DAN | 1.00 | 1.77 | 0.77 | 2.76 | 25 | .011 | 0.52<br>[0.12, 0.92] | E | .032 | Yes |
| LSD (LAM) | VAN | 1.35 | 1.23 | -0.12 | -0.34 | 25 | .740 | -0.06 [-0.44, 0.31] | E | .784 | No |
| LSD (LAM) | LIM | 2.35 | 2.46 | 0.12 | 0.28 | 25 | .784 | 0.05<br>[-0.32, 0.43] | E | .784 | No |
| LSD (LAM) | VIS | 16.65 | 11.08 | -5.58 | -5.37 | 25 | < .001 | -1.02 [-1.48, -0.55] | E | < .001 | Yes |
| LSD (LAM) | SMN | 2.69 | 2.23 | -0.46 | -2.48 | 25 | .020 | -0.47 [-0.86, -0.07] | E | .040 | Yes |
| LSD (LAM) | SUB | 8.81 | 9.77 | 0.96 | 0.82 | 25 | .418 | 0.16 [-0.22, 0.53] | E | .627 | No |
| LSD (LPM) | DMN | 53.97 | 55.43 | 1.47 | 2.26 | 29 | .032 | 0.40 [0.03, 0.76] | P | .032 | Yes |
| LSD (LPM) | FPN | 14.30 | 18.90 | 4.60 | 5.00 | 29 | < .001 | 0.89 [0.47, 1.30] | P | < .001 | Yes |
| LSD (LPM) | DAN | 0.83 | 2.03 | 1.20 | 4.21 | 29 | < .001 | 0.75 [0.35, 1.14] | E | .001 | Yes |
| LSD (LPM) | VAN | 1.60 | 1.27 | -0.33 | -0.93 | 29 | .362 | -0.16 [-0.51, 0.19] | E | .362 | No |
| LSD (LPM) | LIM | 3.83 | 2.60 | -1.23 | -3.22 | 29 | .003 | -0.57 [-0.95, -0.19] | E | .009 | Yes |
| LSD (LPM) | VIS | 13.93 | 11.57 | -2.37 | -2.66 | 29 | .013 | -0.47 [-0.84, -0.10] | E | .025 | Yes |
| LSD (LPM) | SMN | 2.70 | 2.40 | -0.30 | -1.96 | 29 | .059 | -0.35 [-0.71, 0.01] | E | .089 | No |
| LSD (LPM) | SUB | 11.97 | 10.27 | -1.70 | -1.53 | 29 | .138 | -0.27 [-0.62, 0.09] | E | .165 | No |
| PSI | DMN | 54.03 | 55.16 | 1.13 | 1.69 | 30 | .102 | 0.30 [-0.06, 0.64] | P | .102 | No |
| PSI | FPN | 14.26 | 18.77 | 4.52 | 4.61 | 30 | < .001 | 0.81 [0.40, 1.20] | P | < .001 | Yes |
| PSI | DAN | 0.84 | 1.42 | 0.58 | 2.57 | 30 | .015 | 0.45 [0.08, 0.81] | E | .031 | Yes |
| PSI | VAN | 1.58 | 0.87 | -0.71 | -3.05 | 30 | .005 | -0.53 [-0.90, -0.16] | E | .029 | Yes |
| PSI | LIM | 3.97 | 3.16 | -0.81 | -2.17 | 30 | .038 | -0.38 [-0.73, -0.02] | E | .057 | No |
| PSI | VIS | 14.10 | 12.42 | -1.68 | -1.91 | 30 | .066 | -0.33 [-0.68, 0.02] | E | .066 | No |
| PSI | SMN | 2.68 | 2.29 | -0.39 | -2.68 | 30 | .012 | -0.47 [-0.83, -0.10] | E | .031 | Yes |
| PSI | SUB | 12.00 | 9.77 | -2.23 | -1.94 | 30 | .062 | -0.34 [-0.69, 0.02] | E | .066 | No |
| MES | DMN | 53.94 | 55.74 | 1.81 | 2.20 | 30 | .035 | 0.39 [0.03, 0.74] | P | .035 | Yes |
| MES | FPN | 14.10 | 18.13 | 4.03 | 3.58 | 30 | .001 | 0.63 [0.24, 1.00] | P | .002 | Yes |
| MES | DAN | 0.87 | 1.81 | 0.94 | 3.19 | 30 | .003 | 0.56 [0.18, 0.93] | E | .010 | Yes |
| MES | VAN | 1.55 | 1.32 | -0.23 | -0.57 | 30 | .572 | -0.10 [-0.44, 0.24] | E | .572 | No |
| MES | LIM | 4.00 | 2.71 | -1.29 | -3.37 | 30 | .002 | -0.59 [-0.96, -0.21] | E | .010 | Yes |
| MES | VIS | 14.10 | 13.19 | -0.90 | -1.15 | 30 | .258 | -0.20 [-0.55, 0.15] | E | .387 | No |
| MES | SMN | 2.68 | 2.55 | -0.13 | -0.85 | 30 | .403 | -0.15 [-0.49, 0.20] | E | .483 | No |
| MES | SUB | 12.23 | 9.94 | -2.29 | -1.58 | 30 | .124 | -0.28 [-0.63, 0.08] | E | .247 | No |
| MDMA | DMN | 54.88 | 55.65 | 0.77 | 1.18 | 25 | .248 | 0.23 [-0.15, 0.60] | P | .248 | No |
| MDMA | FPN | 15.58 | 17.19 | 1.62 | 1.56 | 25 | .131 | 0.30 [-0.09, 0.68] | P | .248 | No |
| MDMA | DAN | 1.08 | 2.04 | 0.96 | 2.54 | 25 | .018 | 0.48 [0.08, 0.88] | E | .035 | Yes |
| MDMA | VAN | 1.54 | 0.88 | -0.65 | -2.12 | 25 | .044 | -0.40 [-0.79, -0.01] | E | .066 | No |
| MDMA | LIM | 2.42 | 2.27 | -0.15 | -0.51 | 25 | .615 | -0.10 [-0.47, 0.28] | E | .738 | No |
| MDMA | VIS | 16.77 | 13.96 | -2.81 | -3.02 | 25 | .006 | -0.57 [-0.97, -0.16] | E | .017 | Yes |
| MDMA | SMN | 2.69 | 1.88 | -0.81 | -3.89 | 25 | < .001 | -0.74 [-1.16, -0.31] | E | .004 | Yes |
| MDMA | SUB | 8.81 | 8.42 | -0.38 | -0.24 | 25 | .809 | -0.05 [-0.42, 0.33] | E | .809 | No |
| AMPH | DMN | 54.88 | 55.00 | 0.12 | 0.17 | 25 | .869 | 0.03 [-0.34, 0.40] | P | .869 | No |
| AMPH | FPN | 15.77 | 15.46 | -0.31 | -0.36 | 25 | .719 | -0.07 [-0.44, 0.30] | P | .869 | No |
| AMPH | DAN | 1.00 | 1.27 | 0.27 | 1.23 | 25 | .230 | 0.23 [-0.15, 0.61] | E | .276 | No |
| AMPH | VAN | 1.46 | 0.96 | -0.50 | -1.40 | 25 | .173 | -0.27 [-0.64, 0.12] | E | .259 | No |
| AMPH | LIM | 2.46 | 3.23 | 0.77 | 2.18 | 25 | .039 | 0.42 [0.02, 0.80] | E | .077 | No |
| AMPH | VIS | 16.77 | 13.77 | -3.00 | -3.19 | 25 | .004 | -0.61 [-1.01, -0.19] | E | .011 | Yes |
| AMPH | SMN | 2.69 | 1.92 | -0.77 | -3.95 | 25 | < .001 | -0.75 [-1.17, -0.32] | E | .003 | Yes |
| AMPH | SUB | 8.62 | 8.35 | -0.27 | -0.25 | 25 | .802 | -0.05 [-0.42, 0.33] | E | .802 | No |

**Supplementary Table 3.** Limited effect of head motion on changes in gateways per network, per drug condition. ‘Δ raw’ shows the difference in the number of regions that were classified as gateways in each network and drug condition. ‘*q* raw’ indicates the FDR-corrected *p*-value for the gateway-count change in the corresponding network; these values match the corresponding entries in the ‘*q* (tier)’ column in **Supplementary** Table 2. Then, the effect of each drug condition on gateway counts was re-estimated with change in framewise displacement (ΔFD) partialed out (i.e., ΔFD is set to 0). The corresponding adjusted Δ and *q* values are shown. Bolded rows indicate network-drug condition pairs whose significance level changed after the adjustment. There were only two such pairs; the change in DMN gateways in LSD (LPM) lost significance, while the change in LIM gateways in psilocybin gained significance. Correlations between ΔFD and gateway-count changes are also shown, with FDR correction across the 48 condition-by-network correlations. None of these correlations survived multiple comparisons (*q* correlation). Abbreviations: VIS = visual network; SMN = somatomotor network; DAN = dorsal attention network; VAN = ventral attention network; LIM = limbic network; FPN = frontoparietal control network; DMN = default mode network; SUB = subcortical; AMPH = D-amphetamine; PSI = psilocybin; MES = mescaline.

| Drug condition | Network | n | $\Delta$ raw | $q$ raw | $\Delta$ adjusted | $q$ adjusted | $r(\Delta FD, \Delta)$ | $q$ correlation |
| --- | --- | --- | --- | --- | --- | --- | --- | --- |
| LSD (LAM) | DMN | 26 | 0.885 | .196 | 0.921 | .192 | -.076 | .859 |
| LSD (LAM) | FPN | 26 | 4.35 | < .001 | 4.4 | < .001 | -.075 | .859 |
| LSD (LAM) | DAN | 26 | 0.769 | .032 | 0.806 | .027 | -.181 | .753 |
| LSD (LAM) | VAN | 26 | -0.115 | .784 | -0.228 | .573 | .454 | .319 |
| LSD (LAM) | LIM | 26 | 0.115 | .784 | 0.162 | .706 | -.155 | .859 |
| LSD (LAM) | VIS | 26 | -5.58 | < .001 | -5.57 | < .001 | -.012 | .954 |
| LSD (LAM) | SMN | 26 | -0.462 | .040 | -0.474 | .041 | .094 | .859 |
| LSD (LAM) | SUB | 26 | 0.962 | .627 | 0.864 | .573 | .116 | .859 |
| AMPH | DMN | 26 | 0.115 | .869 | -0.078 | .913 | -.224 | .674 |
| AMPH | FPN | 26 | -0.308 | .869 | -0.364 | .913 | -.053 | .886 |
| AMPH | DAN | 26 | 0.269 | .276 | 0.406 | .079 | .5 | .249 |
| AMPH | VAN | 26 | -0.5 | .259 | -0.28 | .479 | .494 | .249 |
| AMPH | LIM | 26 | 0.769 | .077 | 0.791 | .079 | .049 | .886 |
| AMPH | VIS | 26 | -3 | .011 | -2.95 | .019 | .041 | .900 |
| AMPH | SMN | 26 | -0.769 | .003 | -0.74 | .008 | .12 | .859 |
| AMPH | SUB | 26 | -0.269 | .802 | -0.073 | .948 | .147 | .859 |
| MDMA | DMN | 26 | 0.769 | .248 | 0.614 | .347 | -.304 | .674 |
| MDMA | FPN | 26 | 1.62 | .248 | 1.59 | .298 | -.028 | .931 |
| MDMA | DAN | 26 | 0.962 | .035 | 0.986 | .036 | .084 | .859 |
| MDMA | VAN | 26 | -0.654 | .066 | -0.553 | .103 | .419 | .387 |
| MDMA | LIM | 26 | -0.154 | .738 | -0.214 | .583 | -.253 | .674 |
| MDMA | VIS | 26 | -2.81 | .017 | -2.66 | .028 | .202 | .674 |
| MDMA | SMN | 26 | -0.808 | .004 | -0.844 | .003 | -.222 | .674 |
| MDMA | SUB | 26 | -0.385 | .809 | -0.292 | .859 | .075 | .859 |
| **LSD (LPM)** | **DMN** | **30** | **1.47** | **.032** | **1.3** | **.074** | **.13** | **.859** |
| LSD (LPM) | FPN | 30 | 4.6 | < .001 | 4.21 | < .001 | .209 | .674 |
| LSD (LPM) | DAN | 30 | 1.2 | .001 | 1.07 | .008 | .222 | .674 |
| LSD (LPM) | VAN | 30 | -0.333 | .362 | -0.481 | .262 | .205 | .674 |
| LSD (LPM) | LIM | 30 | -1.23 | .009 | -0.946 | .041 | -.374 | .387 |
| LSD (LPM) | VIS | 30 | -2.37 | .025 | -2.71 | .024 | .191 | .674 |
| LSD (LPM) | SMN | 30 | -0.3 | .089 | -0.236 | .236 | -.21 | .674 |
| LSD (LPM) | SUB | 30 | -1.7 | .165 | -1.09 | .357 | -.274 | .674 |
| PSI | DMN | 31 | 1.13 | .102 | 1.27 | .072 | -.185 | .674 |
| PSI | FPN | 31 | 4.52 | < .001 | 4.61 | < .001 | -.085 | .859 |
| PSI | DAN | 31 | 0.581 | .031 | 0.51 | .048 | .264 | .674 |
| PSI | VAN | 31 | -0.71 | .029 | -0.763 | .019 | .194 | .674 |
| **PSI** | **LIM** | **31** | **-0.806** | **.057** | **-0.963** | **.038** | **.357** | **.387** |
| PSI | VIS | 31 | -1.68 | .066 | -1.73 | .075 | .048 | .886 |
| PSI | SMN | 31 | -0.387 | .031 | -0.369 | .039 | -.105 | .859 |
| PSI | SUB | 31 | -2.23 | .066 | -2.21 | .075 | -.013 | .954 |
| MES | DMN | 31 | 1.81 | .035 | 1.76 | .044 | .072 | .859 |
| MES | FPN | 31 | 4.03 | .002 | 4.1 | .003 | -.08 | .859 |
| MES | DAN | 31 | 0.935 | .010 | 0.952 | .011 | -.075 | .859 |
| MES | VAN | 31 | -0.226 | .572 | -0.255 | .532 | .101 | .859 |
| MES | LIM | 31 | -1.29 | .010 | -1.27 | .011 | -.062 | .868 |
| MES | VIS | 31 | -0.903 | .387 | -1.1 | .244 | .345 | .395 |
| MES | SMN | 31 | -0.129 | .483 | -0.108 | .532 | -.192 | .674 |
| MES | SUB | 31 | -2.29 | .247 | -2.08 | .244 | -.193 | .674 |

**Supplementary Table 4.** Motion sensitivity of mean synergy and redundancy PC effects. The raw columns report the two-sided one-sample *t*-test of each within-subject drug-minus-control change in synergy and redundancy PC, averaged across the 235 workspace regions. The adjusted columns report the drug effect after regressing the participation-coefficient change on the corresponding change in framewise displacement (FD). The significance status of each PC effect was unchanged after adjustment. Raw and motion-adjusted *p*-values were each FDR-corrected across the six drug conditions separately for synergy PC and redundancy PC. The correlation column tests the association between ΔFD and the participation-coefficient change; its *q*-value was corrected across all 12 condition-by-measure correlations. None of the correlations were significant after correction.

| Drug condition | Measure | n | *t* raw | *q* raw | *r*(ΔFD, $\Delta$) | *q* correlation | *t* adjusted | *q* adjusted |
| --- | --- | --- | --- | --- | --- | --- | --- | --- |
| LSD (LAM) | Syn PC | 26 | 1.5 | .217 | -.17 | .479 | 1.60 | .183 |
| LSD (LAM) | Red PC | 26 | 7.28 | < .001 | .14 | .548 | 7.03 | < .001 |
| AMPH | Syn PC | 26 | 0.39 | .697 | .26 | .357 | 0.70 | .490 |
| AMPH | Red PC | 26 | 5.95 | < .001 | .44 | .221 | 6.88 | < .001 |
| MDMA | Syn PC | 26 | 0.88 | .464 | .22 | .429 | 1.04 | .369 |
| MDMA | Red PC | 26 | 3.49 | .002 | .04 | .840 | 3.41 | .002 |
| LSD (LPM) | Syn PC | 30 | 3.95 | .001 | .29 | .287 | 3.24 | .008 |
| LSD (LPM) | Red PC | 30 | 11.5 | < .001 | .27 | .287 | 10.52 | < .001 |
| PSI | Syn PC | 31 | 6.02 | < .001 | .34 | .221 | 5.74 | < .001 |
| PSI | Red PC | 31 | 8.82 | < .001 | .33 | .221 | 8.58 | < .001 |
| MES | Syn PC | 31 | 3.29 | .005 | .34 | .221 | 3.15 | .008 |
| MES | Red PC | 31 | 7.33 | < .001 | .18 | .442 | 7.13 | < .001 |

**Supplementary Table 5.** Image acquisition parameters for each of the three fMRI datasets. Abbreviations: HCP = Human Connectome Project; TR = repetition time; TE = echo time; ms = milliseconds; FOV = field of view; L = left; R= right; A = anterior; P = posterior.

|  | HCP | Basel LAM | Basel LPM |
| --- | --- | --- | --- |
| Scanner | 3T Siemens Skyra (customised) | 3T Siemens Prisma | 3T Siemens Prisma |
| Head coil | 32-channel | 20-channel head–neck | 20-channel head–neck |
| Site | HCP consortium | University Hospital Basel | University Hospital Basel |
| Sequence | multiband gradient-echo EPI | 2D gradient-echo EPI | 2D gradient-echo EPI |
| TR | 720 ms | 1,800 ms | 1,800 ms |
| TE | 33.1 ms | 28 ms | 28 ms |
| Flip angle | 52° | 82° | 82° |
| Voxel size | 2.0 mm isotropic | 3.5 mm isotropic, 0.525 mm gap | 3.5 mm isotropic, 0.525 mm gap |
| Slices | 72 | 35 axial, interleaved | 35 axial, interleaved |
| Matrix / FOV | 104 × 90 / 208 × 180 mm | 64 × 64 / 224 × 224 mm | 64 × 64 / 224 × 224 mm |
| Acceleration | multiband 8 | in-plane GRAPPA 2 | in-plane GRAPPA 2 |
| Phase encoding | L→R | A→P | A→P |
| Volumes (duration) | 1,200 (14.4 min) | 300 (9.0 min) | 300 (9.0 min) |
| Anatomical | MPRAGE, 0.7 mm, TR 2,400 / TE 2.14 / TI 1,000 ms, 8° | MPRAGE, 1.0 mm | MPRAGE, 1.0 mm, TE 3.4 ms |

**Supplementary Table 6.** Phenomenology profile of each drug condition. Each entry is the mean placebo-corrected score (drug − placebo) of each drug condition on each of the 11 subscales of the 5D-ASC questionnaire, which was administered in both the LAM and LPM studies.^14,45^ Scores were scaled to the questionnaire maximum before placebo correction. Abbreviations: element = elementary; syn = synesthesia; PSI = psilocybin; MES = mescaline; AMPH = D-amphetamine.

| Drug Cond. | Unity | Spiritual | Bliss | Insight | Dis-embodiment | Impaired control | Anxiety | Complex imagery | Element. imagery | Audio-visual syn. | Changed meaning of percepts |
| --- | --- | --- | --- | --- | --- | --- | --- | --- | --- | --- | --- |
| LSD (LAM) | .485 | .202 | .407 | .336 | .420 | .425 | .149 | .553 | .688 | .657 | .456 |
| LSD (LPM) | .312 | .186 | .326 | .340 | .256 | .237 | .064 | .523 | .547 | .548 | .291 |
| PSI | .235 | .151 | .403 | .311 | .226 | .180 | .055 | .527 | .599 | .484 | .206 |
| MES | .233 | .138 | .286 | .256 | .148 | .127 | .050 | .429 | .466 | .387 | .208 |
| MDMA | .055 | .009 | .222 | .067 | .033 | .027 | .003 | .056 | .047 | .015 | .038 |
| AMPH | .004 | .003 | .087 | .035 | .015 | .008 | .000 | .014 | .000 | .000 | .024 |

